# Phylogeography of *Geothelphusa* freshwater crabs: unexpected dual dispersal routes via land and sea

**DOI:** 10.1101/2022.10.12.511283

**Authors:** Takenaka Masaki, Yano Koki, Tojo Koji

## Abstract

**Aim:** Dispersal is an important factor that determines the potential for colonization to pioneer sites. Most decapods employ seaward migration for reproduction with a planktonic larval phase. However, true freshwater crabs spend their entire life cycle in freshwater. Therefore, it is expected that genetic regionality can be easily detected. In this study, we focused on the genetic structures of true freshwater crabs, *Geothelphusa* crabs. Herein, we reveal the evolutionary history and dispersal patterns of freshwater crustaceans, for which there is limited knowledge to date.

**Location:** Japanese Islands

**Taxon:** *Geothelphusa dehaani* (Decapoda, Potamidae)

**Methods:** We collected and genetically analyzed 283 specimens at 138 localities from freshwater habitats across the Japanese Islands. Phylogenetic analyses were conducted on 1,796 bp of the combined dataset (mtDNA COI, 16S, and nDNA ITS1, histone H3) and 569 bp of the mtDNA COI dataset. The demographic history of *G. dehaani* was simulated using Approximate Bayesian Computation analysis. A salt tolerance experiment was conducted to confirm the survival rate in seawater.

**Results:** The resulting of phylogenetic relationships detected 10 clades that were highly likely to be monophyletic. These 10 clades genetically exhibited an explicit pattern of geographical differentiation. Also, we confirmed the salt tolerance ability of these Japanese freshwater crabs.

**Main conclusions:** The highlights of this study were the discovery of several cryptic species/lineages or undescribed species, and the completely different heterogeneous dual dispersal pathways detected within a single species; i.e., both land and ocean routes. As a result of phylogenetic analysis, it was concluded that Japanese crabs are basically genetically divided by straits. However, strong evidence for dispersion via ocean currents was also detected (i.e., a “sweepstake”), and it was also determined that *G. dehaani* could survive in seawater. This is the first observation of such a unique mode of expansion of a species’ distribution area.

## 1 Introduction

Dispersal is an important factor that determines the potential for colonization to pioneer sites or migration to new habitats, subsequent persistence and the level of genetic connectivity between populations (Banks et al., 2010; García□Verdugo et al., 2017; Van der Stocken et al., 2019; Waters et al., 2020). Population division is also a key process contributing to an organism’s current distribution (Su et al., 1996, 1998; Sota & Vogler, 2001; Tominaga et al., 2013, 2016; Takenaka & Tojo, 2019). The success of colonization to pioneer sites depends on the environmental habitats of the new sites. Phylogeographic patterns influence geographic barriers and the dispersal ability of specific organisms (Avise, 2000; Orsini et al., 2013; Takenaka & Tojo, 2019; Promnun et al., 2021).

Decapoda is a crustacean group diversified in oceans; some members are also adapted to freshwater areas and as well as terrestrial areas. Most decapods employ seaward migration for reproduction, with a planktonic larval phase, even though crabs grow in the freshwater areas of rivers or brackish areas (Kabayashi, 2000; Wowor et al., 2009; Abdullah et al., 2017), i.e., diadromy or catadromy. The distribution patterns and ecology of crabs in riverine environments in Japan has been reported by Kobayashi (2000). The planktonic larval phase(s) predominantly determines potential dispersal ranges (McConaugha, 1992; Rocha et al., 2008; York et al., 2008; Yorisue et al., 2020). Therefore, the geographic genetic structure of most decapods is influenced by ocean currents (Cook et al., 2008; Niikura et al., 2015; Abdullah et al., 2017; Yorisue et al., 2020).

Decapods, including crabs, have evolved to move from the ocean into freshwater zones via brackish water and/or intertidal zones (Pearse, 1927). There are a few species of crab that live in freshwater or terrestrial environments. True freshwater crabs reproduce and spend their entire life cycle in freshwater or terrestrial environments, i.e., without a planktonic larval phase (Kobayashi, 2000; Daniels et al., 2002; Shih et al., 2006, 2011), and they never encounter the ocean (Miyake, 1983), resulting in the relatively poor dispersal abilities of freshwater crabs (Aotsuka et al., 1995; Ikeda et al., 1998). Although adults have the ability for terrestrial movement, freshwater crabs have a strong dependency on riverine habitats (Cook et al., 2008; Daniel et al., 2015). Therefore, it is expected that their genetic structures are affected by river structures and it should be easy to detect genetic regionality (Daniels et al., 2003; Koizumi et al., 2012; Copilaş-Ciocianu et al., 2017).

True freshwater crabs, which have adapted to inland water and also wet terrestrial environments, are distributed globally and there are over 1,300 freshwater species out of 6,700 known species of brachyuran crabs (Yea et al., 2007). These freshwater crabs currently constitute eight families (Pseudothelphusidae, Trichodactylidae, Potamonautidae, Deckeniidae, Platythelphusidae, Potamidae, Gecarcinucidae, and Parathelphusidae: Martin & Davis, 2001; Yea et al., 2007). A freshwater crab taxon, i.e., *Geothelphusa* Stimpson, 1858 (Potamidae), which inhabits mountain streams and/or upper rivers is endemic to Taiwan and the Japanese Islands. In particular, many species are distributed in Taiwan and the Ryukyu Islands (Shokita, 1996; Shokita et al., 2002; Shih et al., 2006, 2007). *Geothelphusa dehaani* is a species endemic to the Japanese Islands, which is widely distributed in Honshu, Shikoku, Kyushu, and surrounding islands (e.g., Yakushima Island, Tanegashima Island, Nakanoshima Island, the Koshikijima Islands, the Oki Islands, Fukuejima Island, Sado Island, and the Kinkazan Island) (Suzuki and Naruse, 2012). On the Osumi Peninsula (Kyushu Island), *Geothelphusa exigua* Suzuki and Tsuda, 1994 is distributed, and *Geothelphusa marmorata* (Suzuki and Okano, 2000) is distributed on Yakushima Island, *Geothelphusa koshikiensis* (Suzuki and Kawai, 2011) is distributed on the Koshikijima Islands, and *Geothelphusa mishima* (Suzuki and Kawai, 2011) is distributed on the Mishima Islands (Fig. 1). In the Ryukyu Islands, different *Geothelphusa* species are distributed on each island.

**Figure 1.**
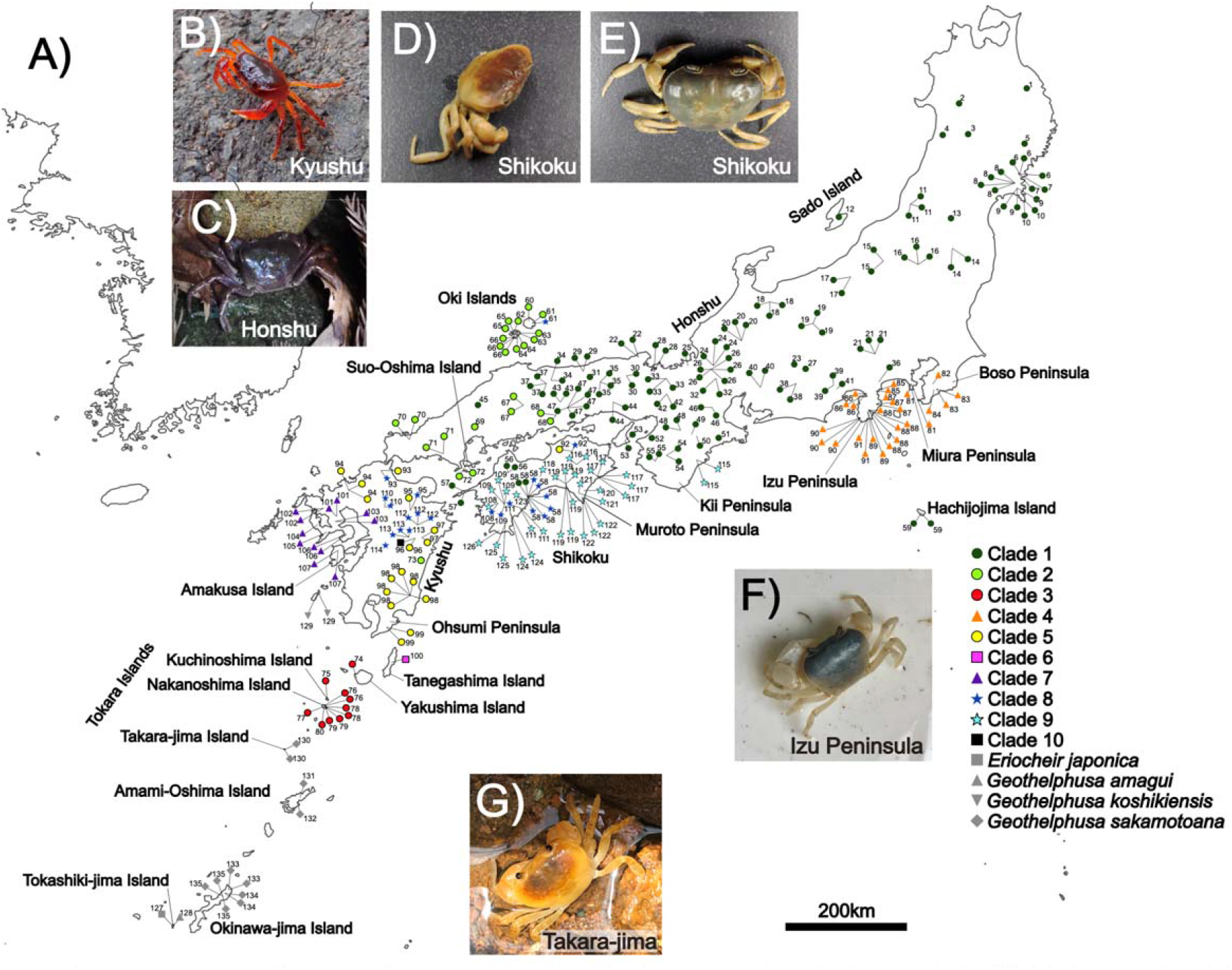
A) Sampling localities and the distribution area of each detected clade (clades 1-10) of the *Geothelphusa* crabs based on data from the mtDNA COI region. Please refer to Table 1 for specific locality numbers, sample numbers, and GenBank accession numbers. The color and number of each clade correspond to the estimated phylogenetic tree shown in Figures 2, 3. B – G) Photos of Sawa crabs on each island. B) Kyushu, C) Honshu, D – E) the same site (st. 116) in Shikoku, F) Izu Peninsula, G) Takara-jima Island.

The habitats of these *Geothelphusa* species are restricted to small streams and ponds with clean freshwater. These habitat characteristics suggest that the vagility or dispersal capability of these crabs is low and led to the genetic differentiation among populations within the species (Nakajima & Masuda, 1985; Ikeda et al., 1998). Regarding *G. dehaani*, populations of Yakushima and Nakanoshima and the Koshikijima Islands have a blue body color, although the body color of the Kyushu Island populations of this species are red (Okano et al., 2000). Chokki (1980) reported that with respect to biogeographical coloration of *G. dehaani*, this species can be differentiated into three color types. Similarly, Suzuki (1992) reported on the regional characteristics of the body color of *G. dehaani* in the North-eastern region of Honshu Island. However, since then all three such body color types have been detected in various regions within Honshu, Shikoku and Kyushu (Chokki, 1976, 1980; Suzuki & Tsuda, 1991; Suzuki, 1992; Furuya and Yamaoka, 2017).

To date, there has been no study of *Geothelphusa* crabs using molecular markers covering their distributional ranges. In this study, we focused on genetic structures at the regional population level and the evolutionary process of true freshwater crabs. These crabs are particularly suitable organisms for discussion regarding how the process of formation of freshwater decapod populations has occurred. Herein, we reveal the evolutionary history of *Geothelphusa* crabs and their genetic regionality, and subsequently compare the genetic structures and dispersal patterns of freshwater crustaceans as observed in some previous studies.

## 2 Materials & Methods

### 2.1 Sampling of materials

Molecular phylogenetic analyses were conducted using 283 specimens including 268 targeted ingroup specimens. Specimens were collected from 138 localities at freshwater habitats across the Japanese Islands between 2015 and 2021. This area reasonably represents almost all of the distribution ranges of *Geothelphusa dehaani* (Toyota, 2019). Almost all the specimens were fixed in 99.5% ethanol in the field, although several specimens were fixed in the laboratory. Detailed information on the specimens used in this study is summarized in Table 1 and Figure 1. Specimens were identified based on the morphological characteristics of the male 1st pleopods (Toyota & Seki, 2019), which are considered to exhibit key identifying characteristics.

**Table 1.**
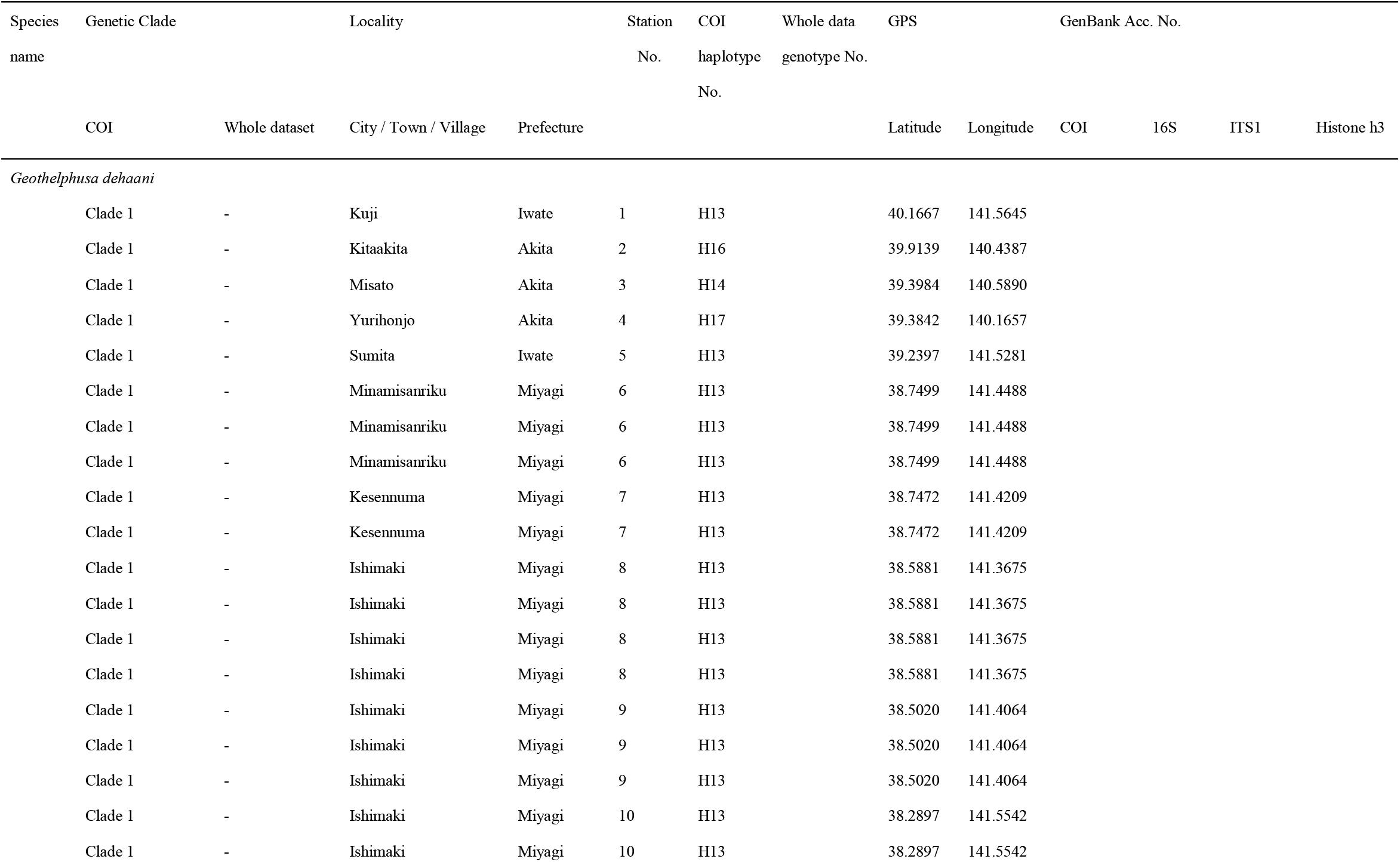
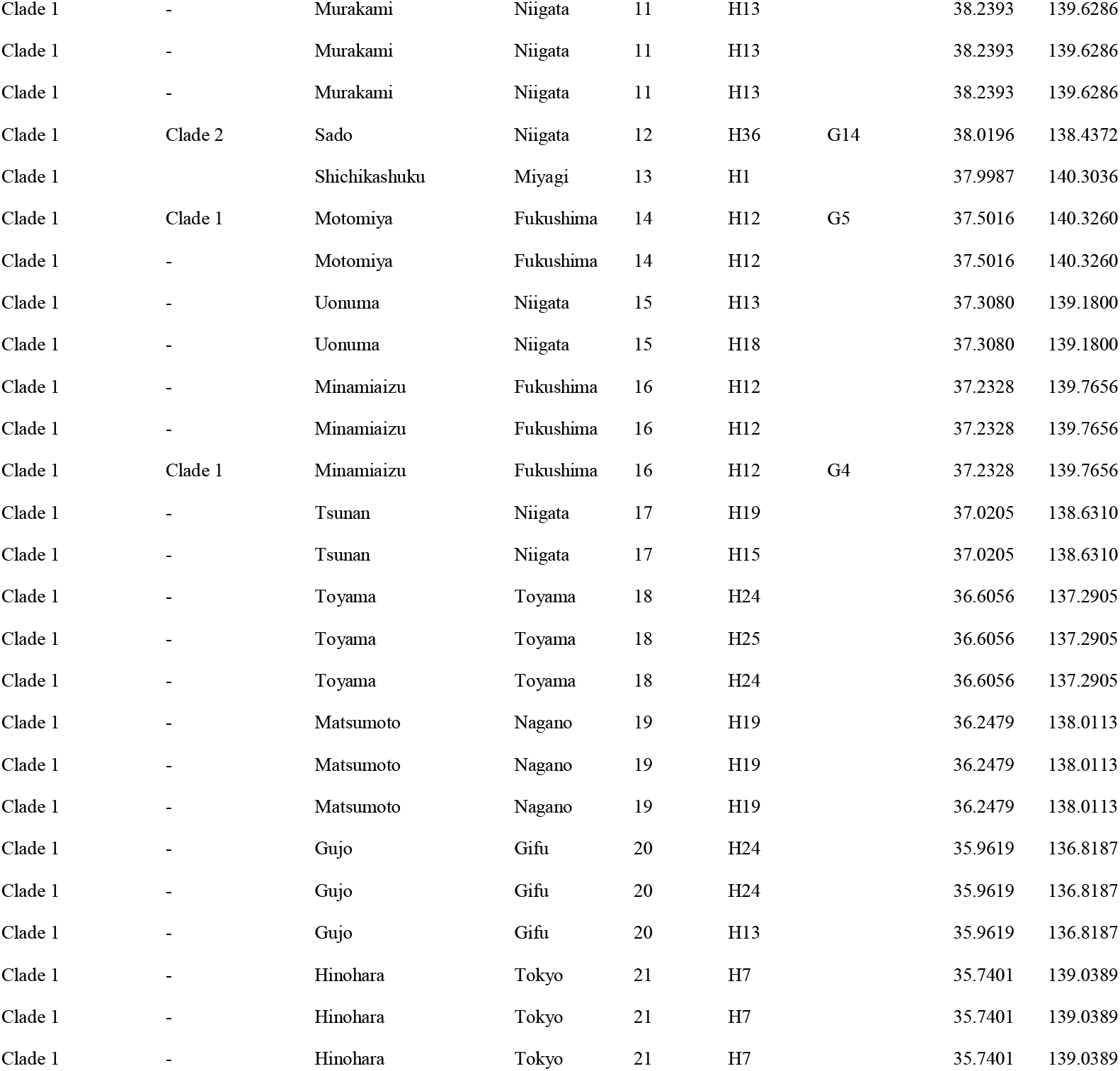
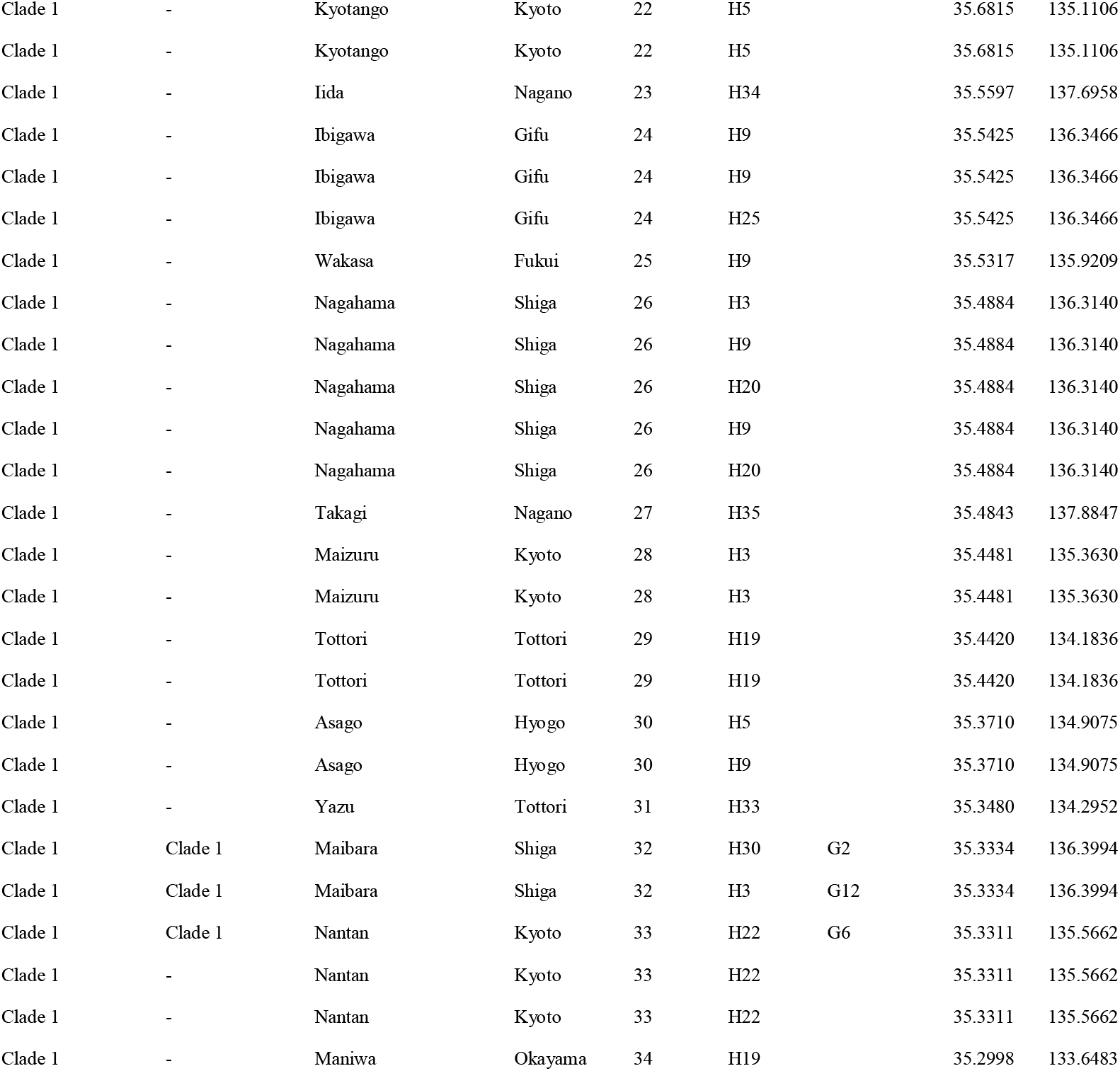
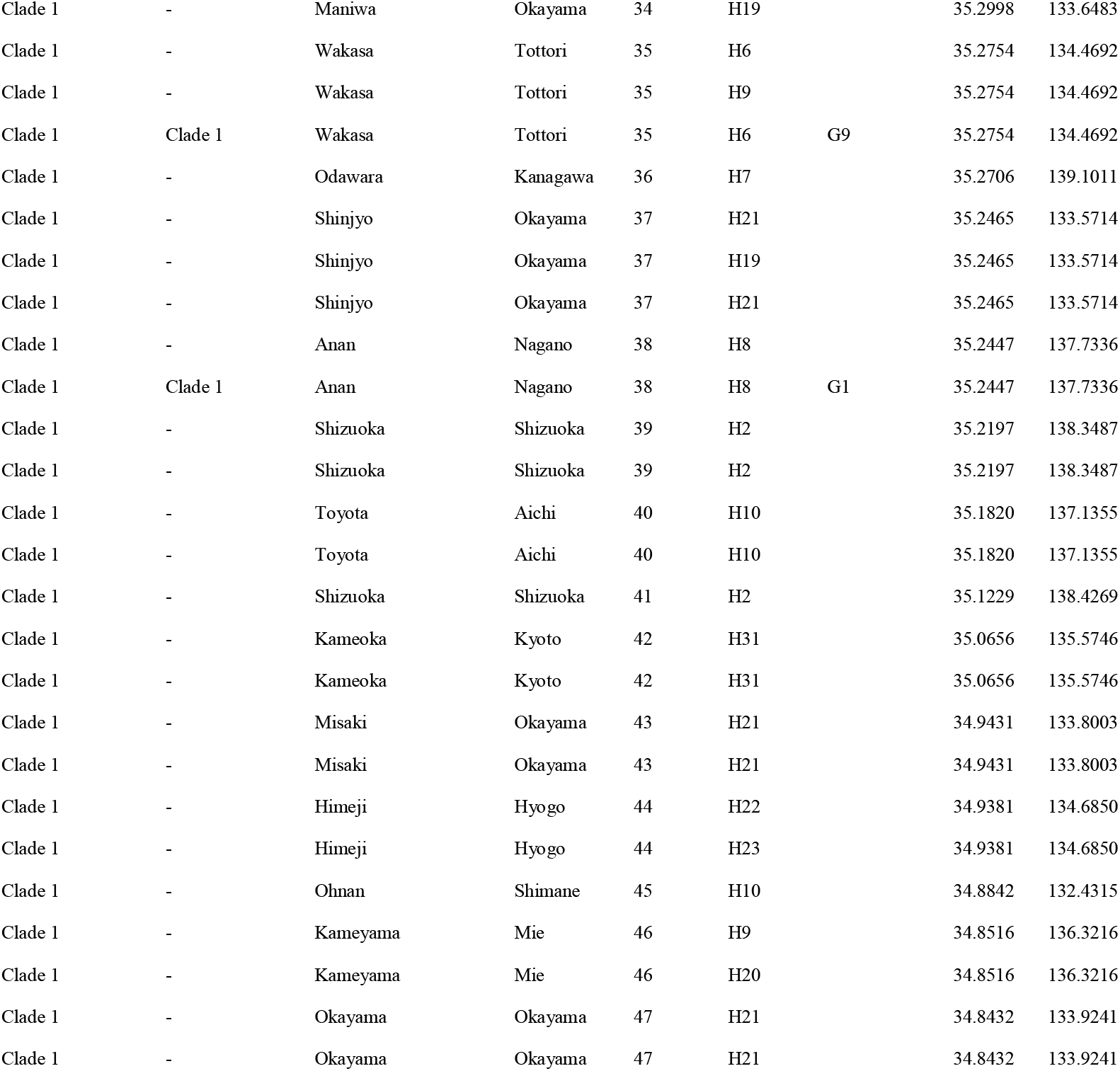
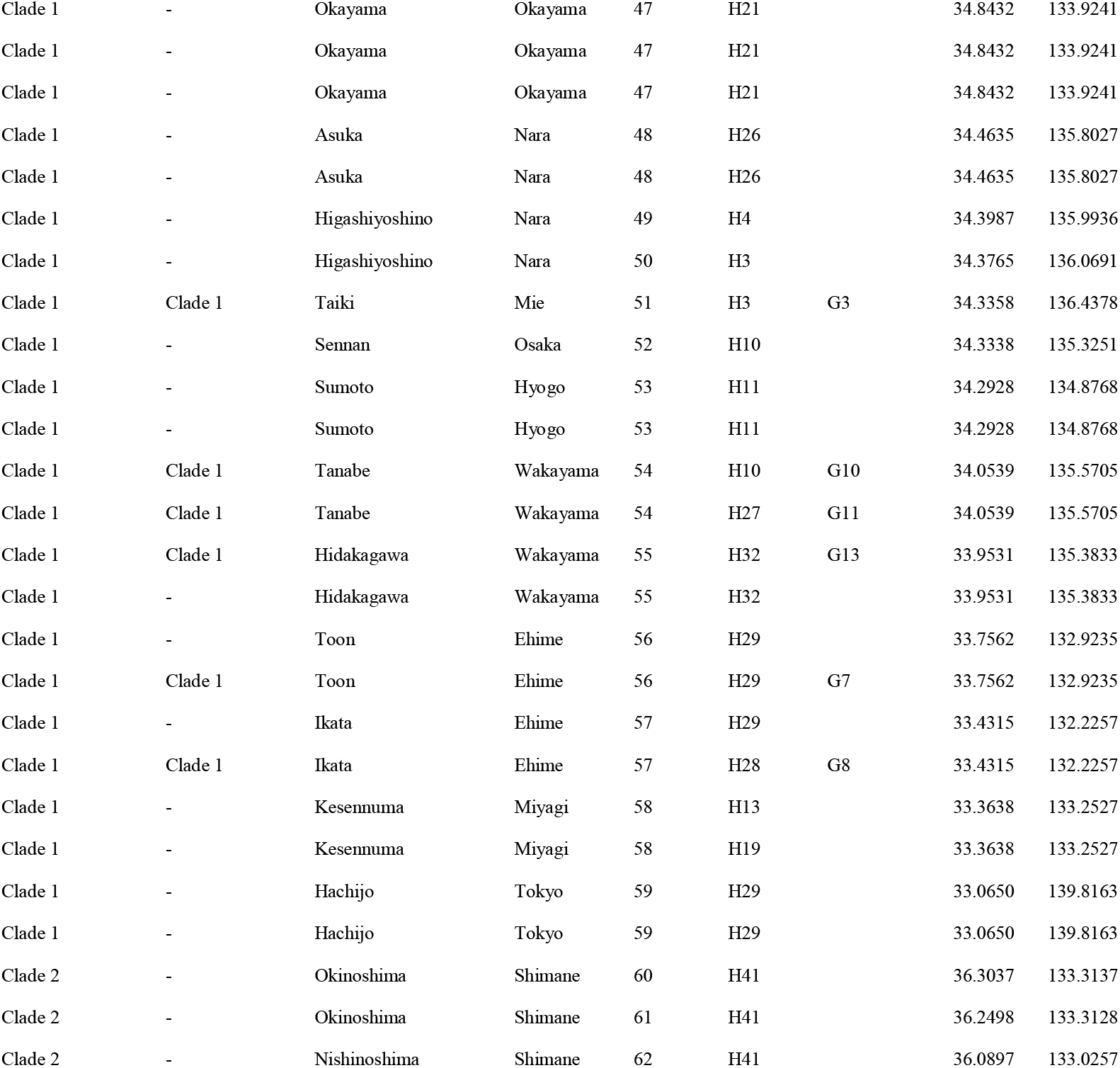
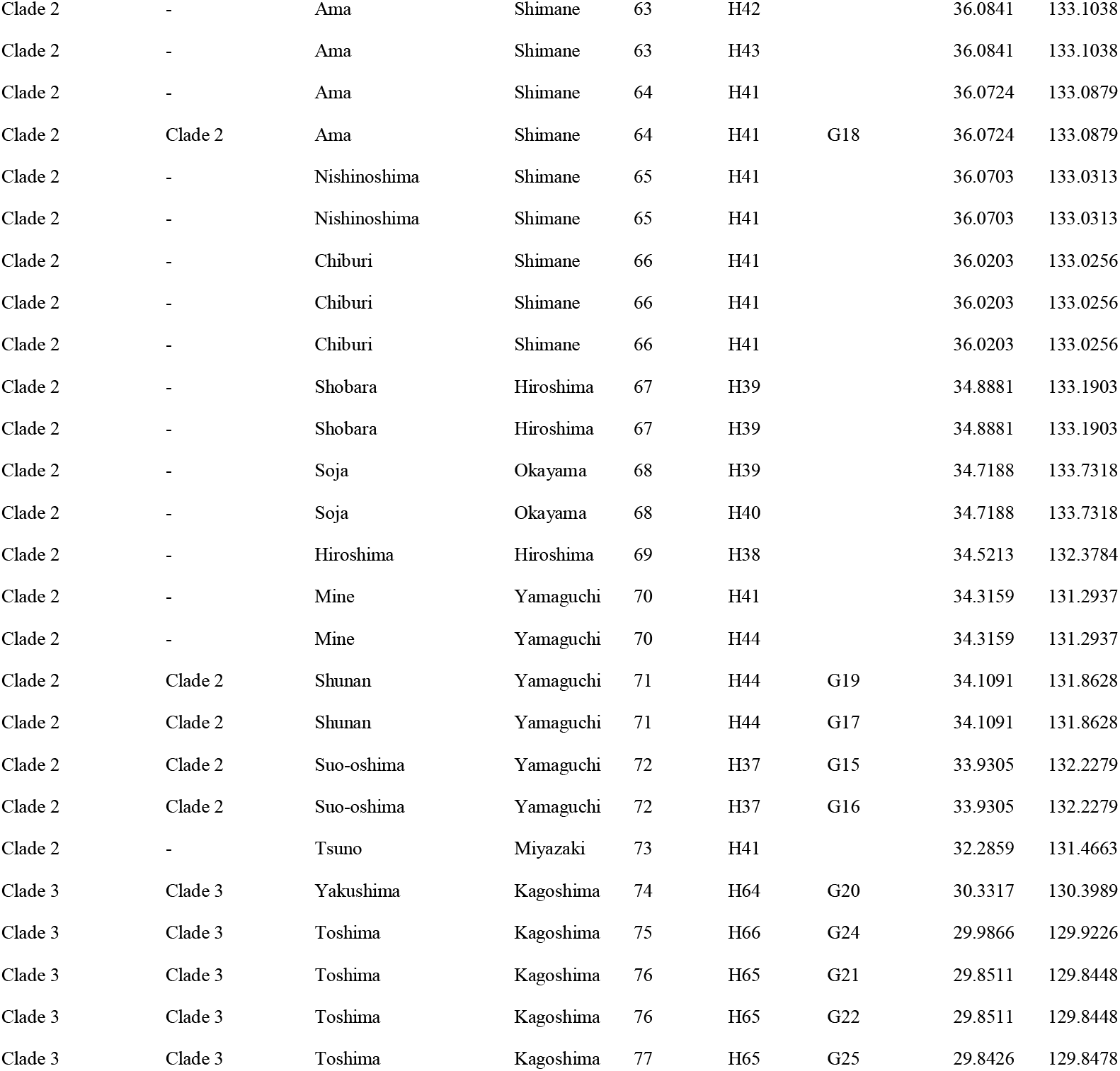
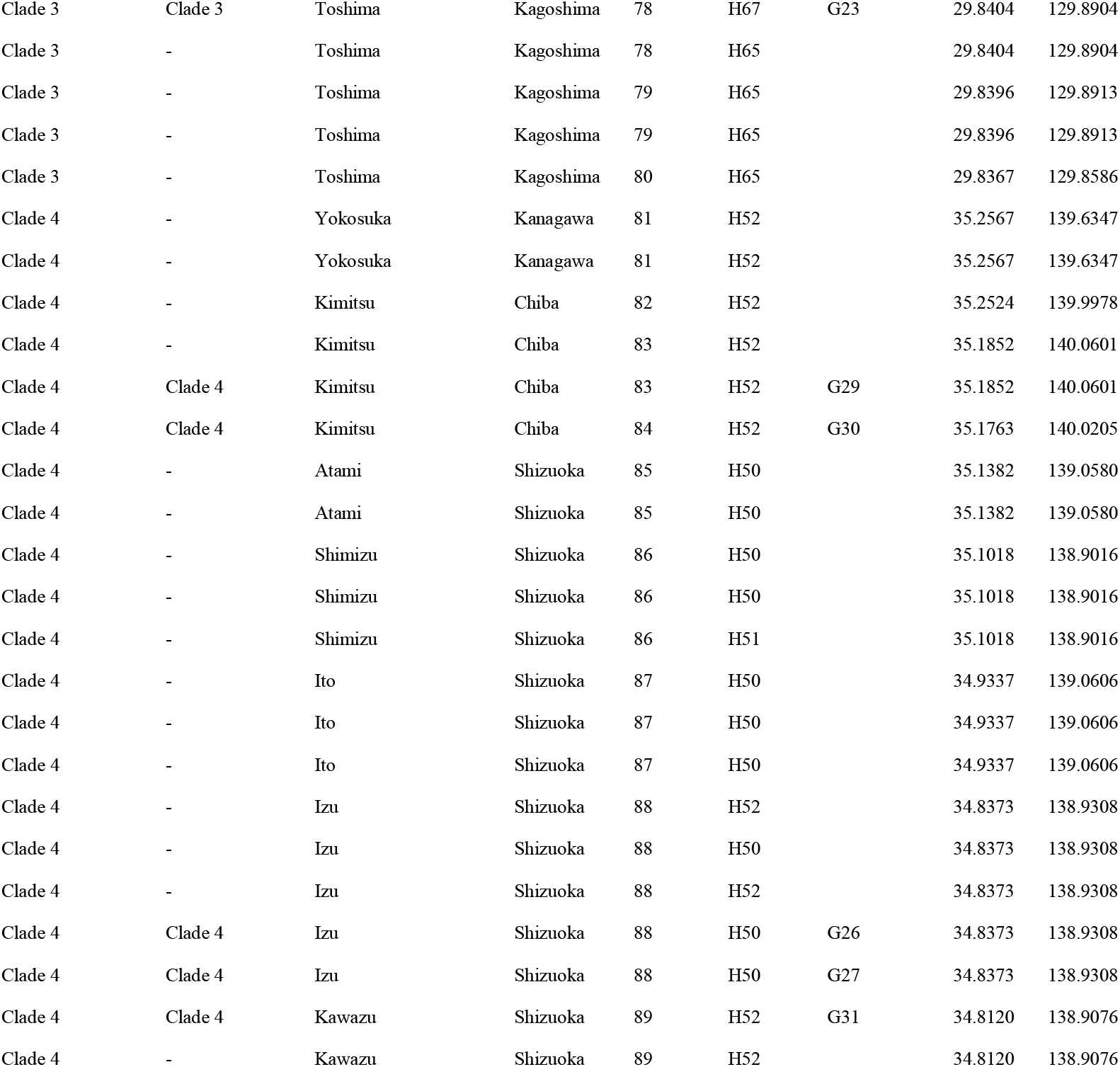
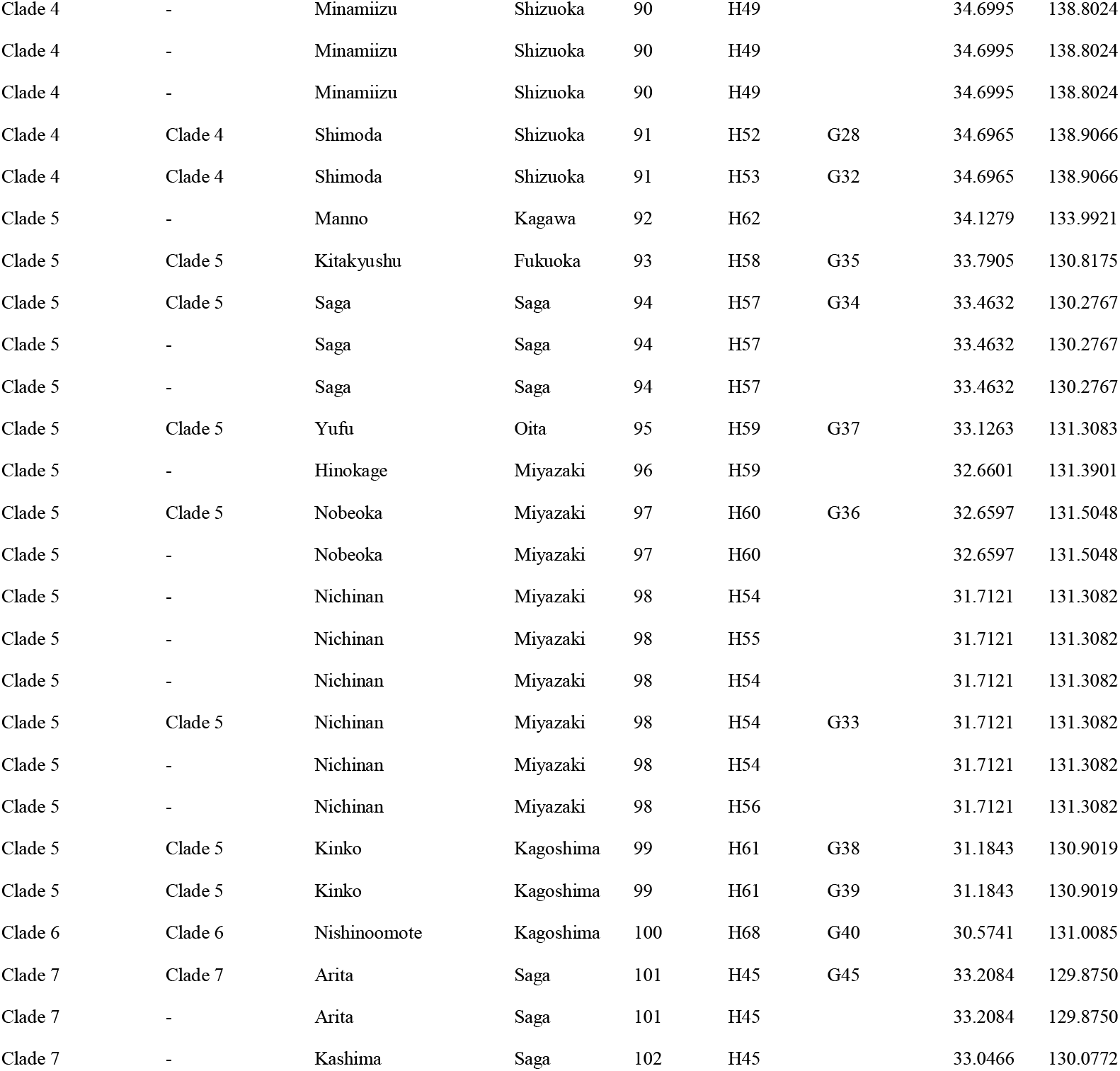
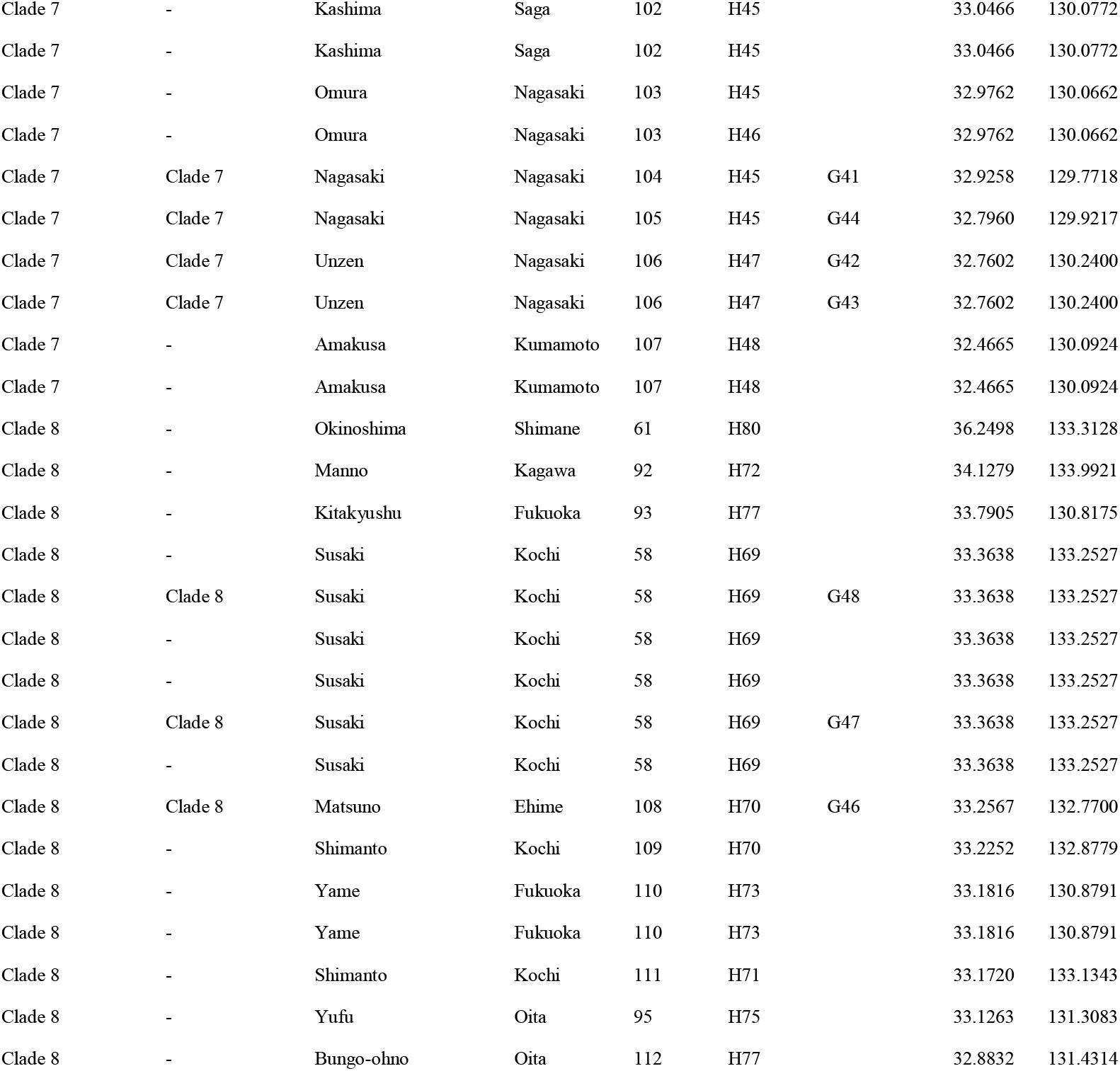
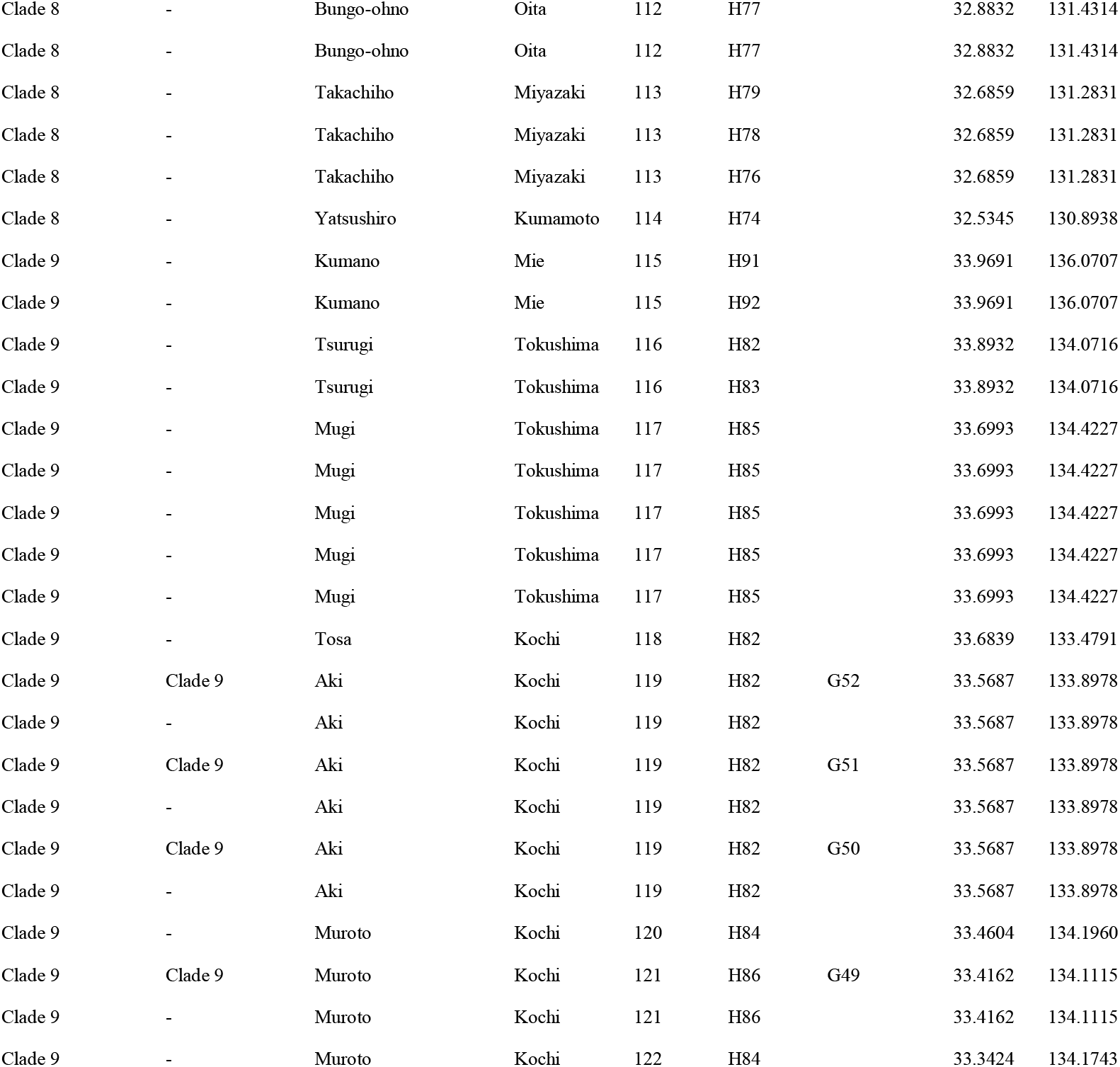
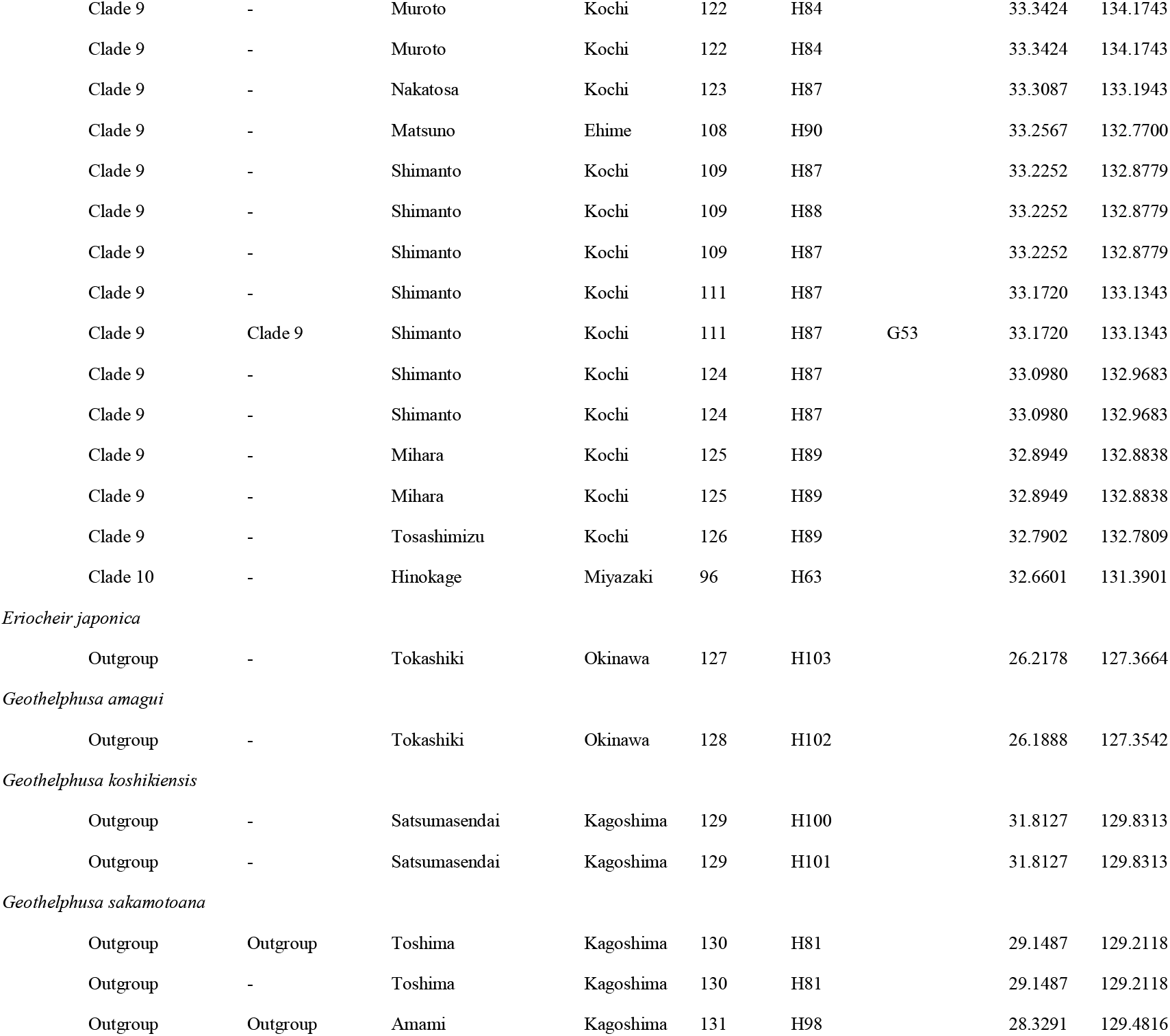
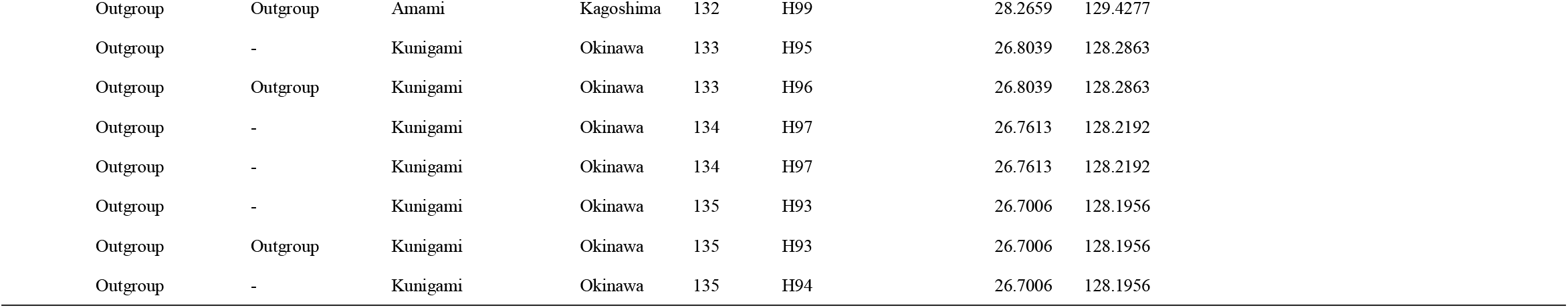
List of specimens of *Geothelphusa* crabs examined in this study, sequence types, and the GenBank accession numbers

### 2.2 DNA extraction, amplification, sequencing and alignment

Total genomic DNA was extracted from a tissue sample from ethanol-preserved specimens and DNA was purified using a DNeasy Blood & Tissue Kit (Qiagen, Hilden), according to the manufacturer’s instructions. Total DNA was used to amplify fragments of mitochondrial DNA (mtDNA) cytochrome c oxidase subunit I (COI), 16S rRNA, nuclear DNA (nDNA) internal transcribed spacer 1 (ITS1) and histone H3. The primer sets used for polymerase chain reaction (PCR) and annealing temperatures are listed in Table 2. rTaq (TOYOBO, Osaka) was used for COI, ExTaq (Takara, Shiga) for 16S, ITS1, and histone H3. PCR products were purified using ExoSAP-IT (GE Healthcare, Buckinghamshire). Purified DNA fragments were used for cycle sequencing reaction with a BigDye Terminator v 1.1 Cycle Sequencing Kit (Applied Biosystems, California) and sequenced on an automated DNA Sequencer (ABI3130xl DNA analyzer; Applied Biosystems, California).

**Table 2.**
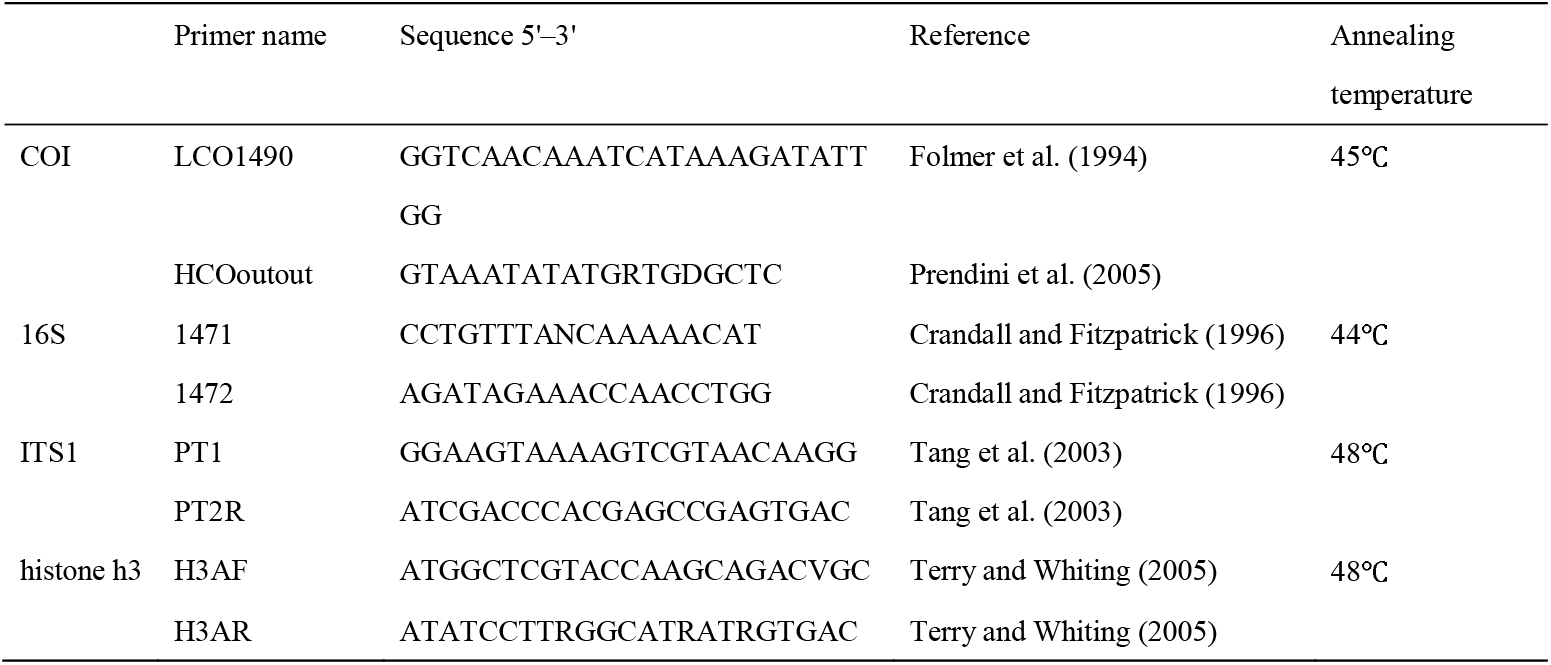
Primers used in this study

All of the sequence data have been submitted to the DNA data bank of Japan (DDBJ database) and accession numbers are provided in Table 1. Sequence alignment and editing were conducted using MEGA ver. 6.06 (Tamura et al., 2013) and CLC working bench (CLC bio, Aarhus). All sequence data were aligned automatically by Clustal W (Thompson et al., 1994) instrumented in MEGA ver. 6.06 (Tamura et al., 2013).

### 2.3 Phylogenetic analysis

Phylogenetic analyses were conducted on 1,796 bp of the combined dataset (mt DNA COI, 16S, and nDNA ITS1, histone H3) and 569 bp of the mtDNA COI dataset, using the maximum likelihood method (ML; Felsenstein, 1981) and Bayesian inference (BI; Huelsenbeck et al., 2001). ML analysis was performed with RAxML (Stamatakis, 2014) using the GTR + G + I model and 1,000 bootstrap replicates. Bayesian inference was conducted using MrBayes5D ver 3.1.2 (Tanabe, 2008) and the HKY85 + G + I model for the combined dataset and the HKY85 + G model for the mtDNA COI dataset, which is the best fit model selected based on the Schwartz’s Bayesian Information Criterion (BIC; Schwartz, 1978) in Kakusan 4 (Tanabe, 2007). Two Markov Monte Carlo chains were run for 10 million generations with sampling every 1000 generations. The first 10% of the sampling trees were discarded as burn-in. Bayesian log trace files were visualized by Tracer ver. 1.6 (Rambaut & Drummond, 2007) and we confirmed the effective sampling size (ESS) was more than 200. Both ML and Bayesian phylogenetic trees were visualized by FigTree ver.1.3.1 (Rambaut, 2009).

### 2.4 Demographic history

Demographic history of *Geothelphusa dehaani* was simulated using a combined dataset. Based on the results of phylogenetic analysis, Clades 1–5 and Clade 7 were clustered in six groups for demographic analysis. Because only single specimens were detected in this study, Clade 6 was excluded from the analysis. To test the phylogenetic relationship and the possibility of oceanic dispersal by the Kuroshio Current in this species, approximate Bayesian computation (ABC) analysis implemented in DIYABC ver. 2.1.0 (Cornuet et al., 2014) were used to compare five scenarios. As shown in Figure 2, the genealogical position of Clade 4 was tested in the analysis. All possible summary statistics were included and 6,000,000 simulations were run for the scenario. The best scenario was selected from the results of direct estimation and the logistic regression estimate of the posterior probabilities for scenarios. All the DIYABC results (summary statistics) are shown in Figure 2.

**Figure 2.**
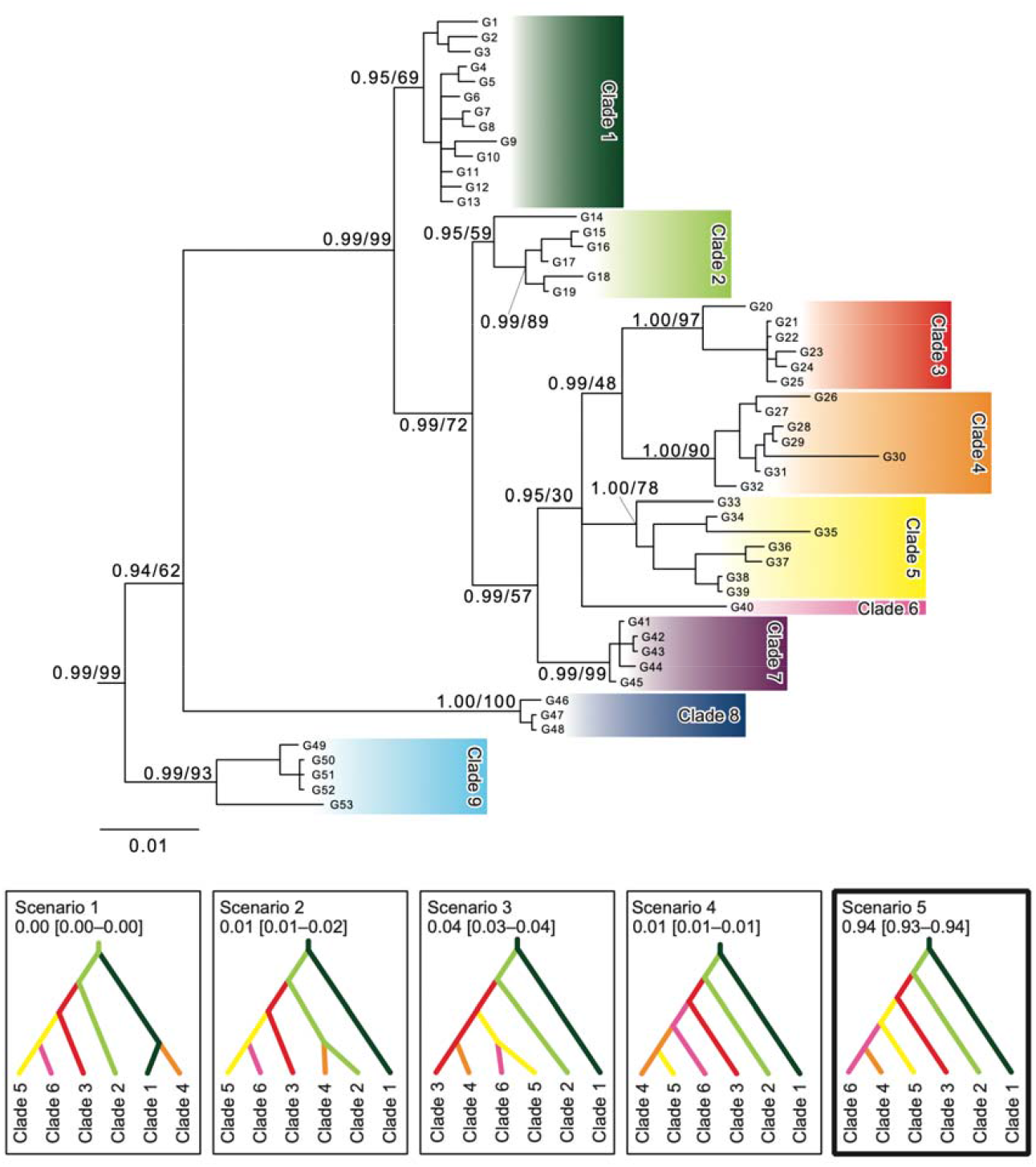
The estimated phylogenetic relationships [Bayes tree] of the *Geothelphusa* crabs based on the sequenced data of the combined dataset (mt DNA COI, 16S, and nDNA ITS1, histone h3: 1,796 bp) using 58 specimens. The number of major nodes indicates posterior probabilities and bootstrap values. The color and number of each clade correspond to the map of sampling localities shown in Figure 1.

### 2.5 Salt Tolerance Evaluation

A salt tolerance experiment was conducted to confirm the survival rate in seawater. We collected *G. dehaani* including adults and juveniles from the Izu Peninsula (St. 88) and Nagano (St. 19) (Table 1), and one more site (Sanada, Ueda, Nagano: 36.4963, 138.3622) not used for genetic analysis. In a laboratory, *G. dehaani* were reared in artificial seawater (Tetra Marine Salt Pro; USA) for 15 days, and their survival was confirmed every day.

As artificial water, 34 g of “Tetra Marine Salt Pro” were added to 1,000 mL tap water (salinity of ca. 3.4%), which is a general seawater concentration, with salinity of 6.8% (Tetra Marine Salt Pro is 68 g in 1,000 mL tap water), which is twice that concentration, were used for the salt tolerance experiment. The rearing container was a Plastic case (NAKAYA KAGAKU SANGYO CO.; Japan; 173×118×118 mm). No food was given during the experiment to avoid its influence; it is unlikely that food will be eaten if a crab disperses by ocean currents (there is no problem in terms of survival even if no food is given for 15 days).

In the experiment, freshwater crabs were reared in three experiments: seawater, twice the concentration of seawater, and freshwater (tap water). After 15 days, the freshwater crabs reared in freshwater were then placed in seawater, and the freshwater crabs reared in seawater were placed in freshwater, respectively. The freshwater crabs reared in twice the concentration of seawater did not survive for 15 days, so the experiment was terminated.

## 3 Results

### 3.1 Phylogenetic analyses

The results of phylogenetic relationships observed for *Geothelphusa dehaani* specimens, based on the mtDNA COI region (569 bp) for 283 specimens, and based on the combined datasets of the mtDNA COI (560 bp) and 16S rRNA (532 bp) regions, and the nDNA ITS (440 bp) and histone H3 (264 bp) regions using 58 specimens, are shown in Figures 3 and 2, respectively. The phylogenetic cladogram based on each data set detected 10 clades, and almost all of the clades were consistent between the COI dataset and the combined total dataset (Table 1). The distribution of the collection sites of the specimens in each clade is shown in Figure 1. These 10 clades were highly likely to be monophyletic (Figs. 2, 3). Genetic relationships between respective clades based on the mtDNA COI region dataset were not observed (Fig. 3), while genetic relationships based on the combined dataset were more clearly observed (Fig. 2). However, the phylogenetic cladogram could not reliably reveal the relationships among clades 3 to 6.

**Figure 3.**
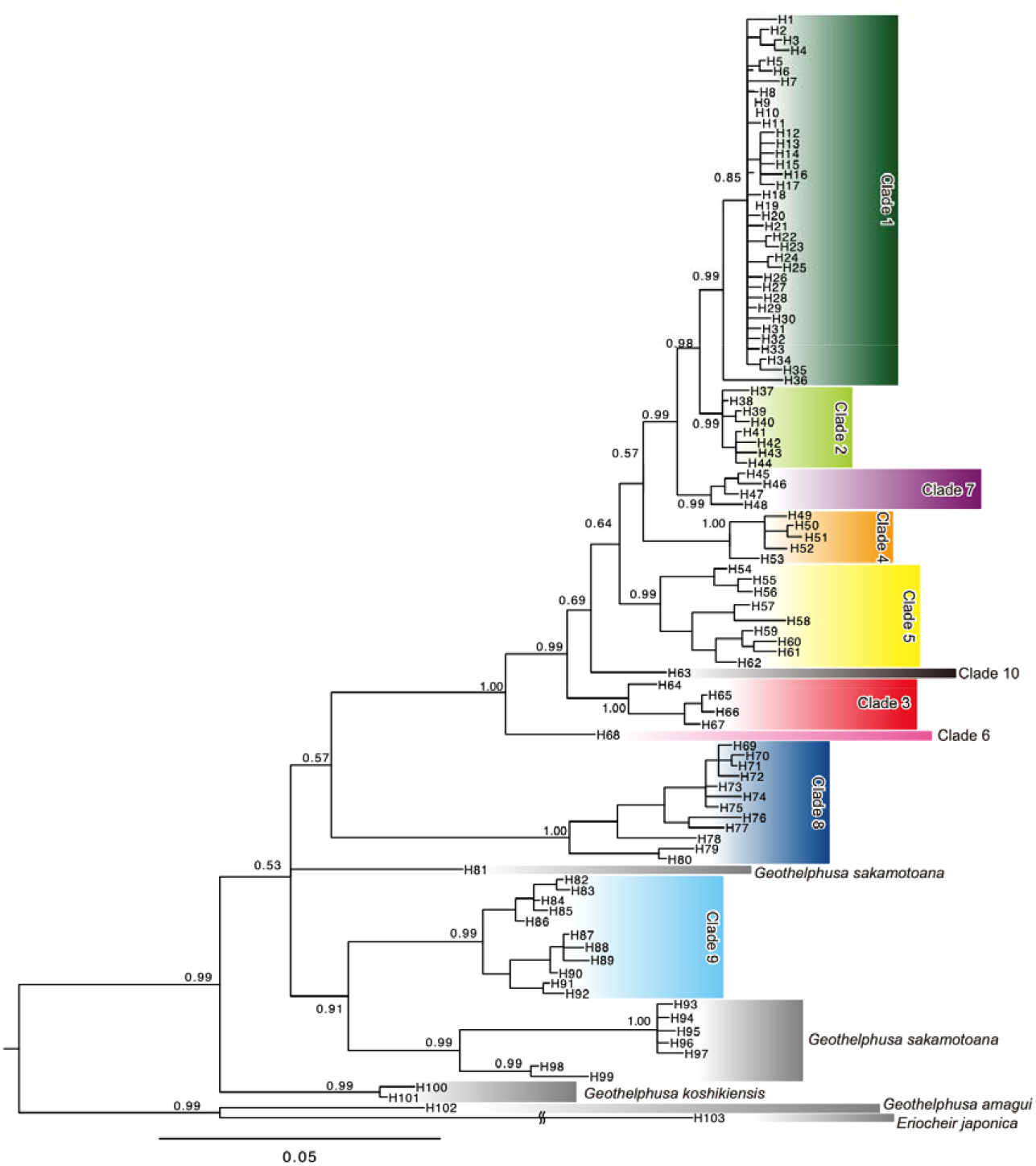
The estimated phylogenetic relationships [Bayes tree] of the *Geothelphusa* crabs based on the sequenced data of the mtDNA COI region (569 bp) using 283 specimens. The number major nodes indicates posterior probabilities. The color and number of each clade correspond to the map of sampling localities shown in Figure 1.

Based on the mtDNA COI region, Clade 1 is mainly distributed in Honshu, with some specimens from Shikoku and Sado Island. Clade 2 is distributed in Western Honshu, and Suo-Oshima Island, the Oki Islands, and a single specimen from Kyushu was included. Clade 3 is distributed on the Tokara Islands and Yakushima Island (i.e., the Northern Ryukyu Islands). Clade 4 is distributed on the Izu Peninsula, the Boso Peninsula, and the Miura Peninsula. Clade 5 is mainly distributed in Kyushu, with one specimen from Shikoku. Clade 6 is distributed on Tanegashima Island. Clade 7 is distributed in Northwestern Kyushu (i.e., the Shimabara Peninsula, and the Amakusa Islands). Clade 8 is distributed in Shikoku and Kyushu. Clade 9 is mainly distributed in Shikoku, and some in Honshu. Clade 10 is distributed in Kyushu. The genetic distances (*p*-distances) between respective clades are shown in Table 3. Clades 8 and 9 had a greater genetic distance from each other than the genetic distances observed between other clades. The genetic diversity (i.e., both the haplotype diversity and nucleotide diversity) of each clade is shown in Table 4. In addition, recent expansion of the effective population sizes of Clade 1 was clearly indicated (see Tajima’s D and Fu’s Fs; Table 4).

**Table 3.** Genetic distance (*p*-distance) between each clade of *Geothelphusa dehaani* based on the mtDNA COI (569-bp) region.

|  | Clade1 | Clade2 | Clade3 | Clade4 | Clade5 | Clade6 | Clade7 | Clade8 |
| --- | --- | --- | --- | --- | --- | --- | --- | --- |
| Clade2 | 0.015 | - | - | - | - | - | - | - |
| Clade3 | 0.041 | 0.041 | - | - | - | - | - | - |
| Clade4 | 0.032 | 0.034 | 0.041 | - | - | - | - | - |
| Clade5 | 0.031 | 0.035 | 0.041 | 0.042 | - | - | - | - |
| Clade6 | 0.048 | 0.049 | 0.046 | 0.051 | 0.046 | - | - | - |
| Clade7 | 0.021 | 0.019 | 0.04 | 0.037 | 0.029 | 0.046 | - | - |
| Clade8 | 0.090 | 0.090 | 0.091 | 0.087 | 0.092 | 0.083 | 0.084 | - |
| Clade9 | 0.075 | 0.079 | 0.082 | 0.079 | 0.081 | 0.082 | 0.074 | 0.077 |

**Table 4.** Mismatch distribution analysis and neutrality tests of each clades

| | n | Hd | $\pi$ | Tajima's D | Fu's Fs |
| --- | --- | --- | --- | --- | --- |
| All samples | 268 | 0.981 | 0.042 | -0.533 | **-17.942\*** |
| Clade1 | 120 | 0.949 | 0.006 | **-1.942\*** | **-33.646\*** |
| Clade2 | 22 | 0.697 | 0.003 | -1.155 | -2.305 |
| Clade3 | 9 | 0.583 | 0.002 | -1.678 | -0.822 |
| Clade4 | 29 | 0.810 | 0.010 | -1.298 | 1.634 |
| Clade5 | 17 | 0.912 | 0.018 | 0.487 | 1.649 |
| Clade6 | 1 | - | - | - | - |
| Clade7 | 13 | 0.615 | 0.004 | -0.599 | 1.283 |
| Clade8 | 22 | 0.900 | 0.018 | -0.895 | 0.492 |
| Clade9 | 34 | 0.877 | 0.012 | 0.963 | 0.324 |

n, sample size; Hd, haplotype diversity; $\pi$ , nucleotide diversity; \*, $p < 0.01$

Regarding the inconsistent results of phylogenetic analyses based on the dataset of only the mtDNA COI region and the combined dataset, specimens from Sado Island (St. 12) were found to have contradicting results (Table 3; Figs. 2, 3). A specimen from Tanegashima Island (St. 100; Clade 6) was analyzed by means of both the COI based dataset and the combined dataset, and was also found to be greatly genetically differentiated from the other clades. A specimen from Kyushu (St. 95; Clade 10) was also found to be greatly genetically differentiated from the other clades based on only the mtDNA COI region.

### 3.2 Demographic history

ABC analysis was conducted in order to reveal the most probable phylogenetic relationships between clades which had less reliability based on phylogenetic analysis results, and to reveal the genealogical position of Clade 4 that mainly consists of specimens from isolated peninsulas. In a comparison of the direct estimation results of calculated posterior probabilities between scenarios, Scenario 2 was the most supported and the second most supported scenario was Scenario 5 [posterior probabilities and confidence intervals of Scenarios 1–5: 0.17 (0.00–0.51); 0.29 (0.00–0.68); 0.17 (0.00–0.50); 0.13 (0.00–0.42); 0.24 (0.00–0.62), respectively]. From the results of the logistic regression estimate for scenarios, Scenario 5 was the most supported [posterior probabilities and confidence intervals of Scenarios 1–5: 0.00 (0.00–0.00); 0.01 (0.01–0.02); 0.04 (0.03–0.04); 0.01 (0.01–0.01); 0.94 (0.93–0.94), respectively].

### 3.3 Salt tolerance evaluation

Since the results of the phylogenetic analysis indicated the probability of ocean current-based dispersion (i.e., the Kuroshio current), we evaluated the salt tolerance ability of *G. dehaani*. As a result, no significant difference in survival was detected between freshwater and seawater species for at least 15 days (Table 5; Fig. 4). However, Clade 4 of *G. dehaani* could not survive when the concentration of the seawater saltiness was doubled (n = 5). We also investigated whether or not they could survive when they arrived in new freshwater environments after dispersal via ocean currents. When they were reared in freshwater after being reared in seawater for 15 days, there was no detrimental effect on their survival. There was no difference in salt tolerance between Clades 1 and 4.

**Table 5.**
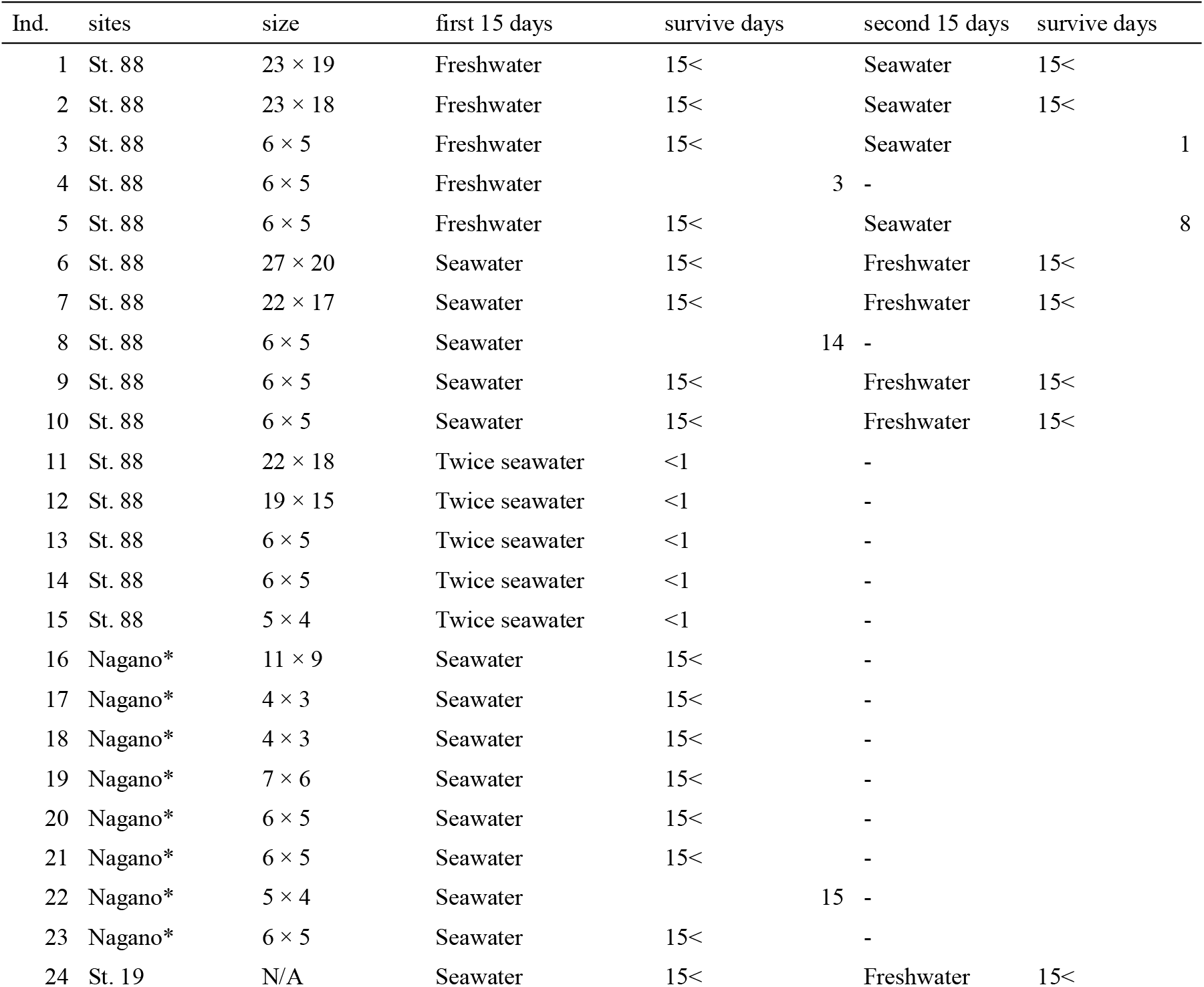
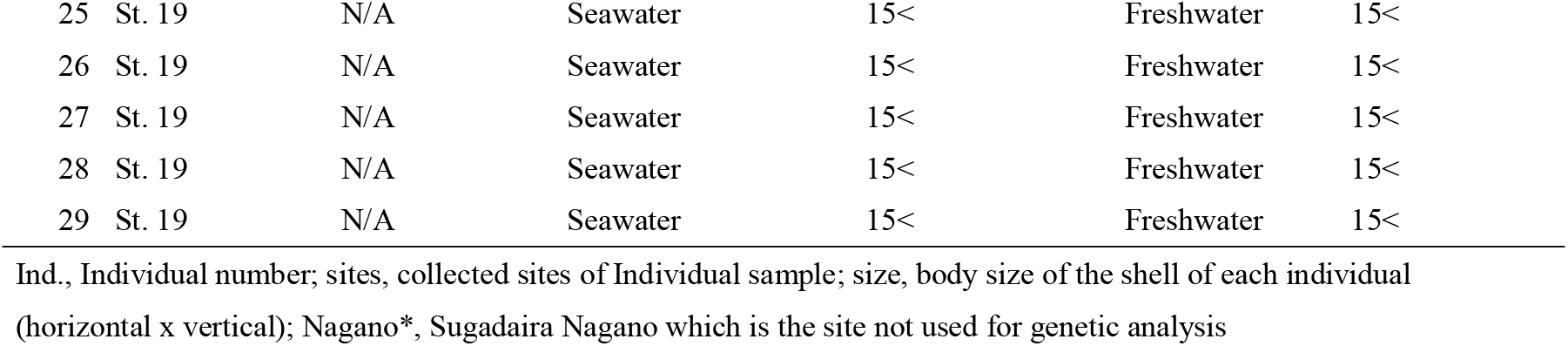
The experiment of the salt tolerance evaluation for *Geothelphusa dehaani*

**Figure 4.**
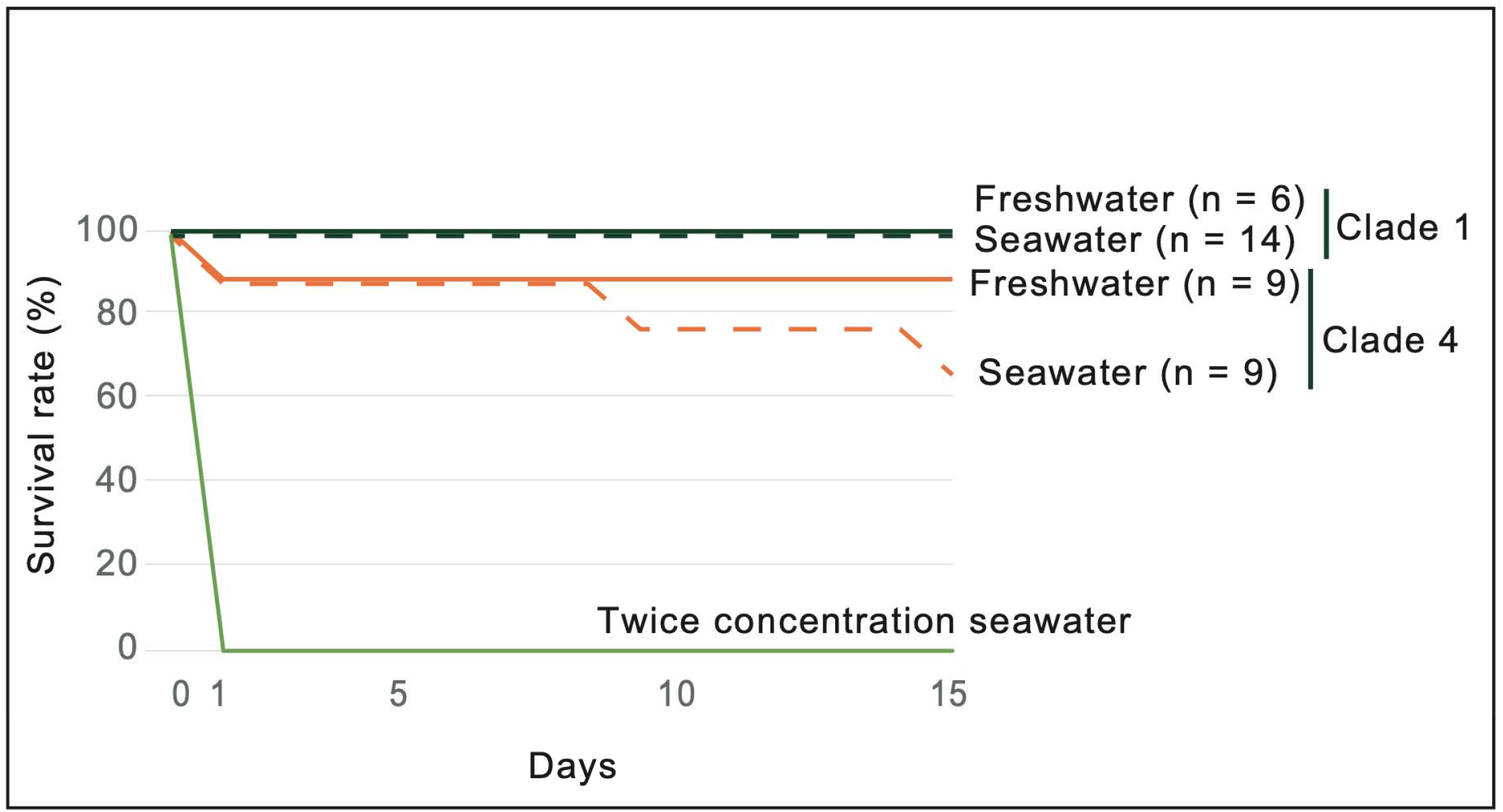
Survival rate of *Geothelphusa dehaani* of Clades 1 and 4 for 15 days in water with different salinity levels (freshwater, seawater, twice concentration seawater). The colors of survival curves correspond to each clade color in Figure 1.

## 4 Discussion

The diversity of the *Geothelphusa* genus is very wide, especially in Southeast Asia and the Ryukyu Islands, so it is thought that the origin of the freshwater crab *Geothelphusa* was in the more southern regions, with subsequent migration northward through the Ryukyu Islands (Shih & Ng, 2011; Shih et al., 2004, 2011). Regarding Honshu, Shikoku, Kyushu, and the surrounding islands, *Geothelphusa exigua* is distributed in the southern part of Kyushu, i.e., the Osumi Peninsula (Suzuki & Tsuda, 1994), *Geothelphusa marmorata* is distributed on Yakushima Island (Suzuki & Okano, 2000), and *Geothelphusa mishima* is distributed on the southwestern islands of Kyushu (i.e. Kuchinoerabu Island, Kuroshima Island; Suzuki & Kawai, 2011), *Geothelphusa koshikiensis* is distributed on the Koshiki Islands (Suzuki & Kawai, 2011). As such, there is a high degree of species diversity in the south western parts of the Japanese Islands. Also, in this study, it was revealed that the four genetic clades (i.e., Clades 5, 7–8 and 10) within the species of *G. dehaani* are distributed in Kyushu, and since the genetic diversity of southwestern Japan is high, it is considered that *G. dehaani* also originated from the southwestern direction and migrated northward.

### 4.1 Genetic differentiation by island

It is known that many crustaceans disperse over a wide area via ocean currents due to their planktonic life cycle during their larval stages in brackish waters and/or the ocean. Therefore, they have significant potential to disperse via ocean currents (McConaugha, 1992; Rocha et al., 2008; York et al., 2008; Abdullah et al., 2017; Yorisue et al., 2020). It is well documented that the distributional ranges of geographical lineages of marine organisms are greatly influenced by ocean currents (Rocha et al., 2008; York et al., 2008). The larvae of decapods are generally dispersed over wide areas by oceanic surface currents (Cook et al., 2008; Page et al., 2008). However, as for the freshwater crab, *G. dehaani*, it spends its entire life in freshwater, so it has no stages in which it can migrate via the ocean. Therefore, *G. dehaani* was expected to be an easy species in which to detect genetic differentiation within each geographical region. As a result, *G. dehaani* was shown clearly to be composed of nine major genetically differentiated clades, based on the sets of mtDNA COI, 16S rRNA, and nDNA histone H3 regions. Genetic differentiation of each clade was primarily detected between islands. This means that each strait functions as a large genetic barrier for this freshwater crab, as was initially expected. Other related species of the same *Geothelphusa* genus were distributed around Kyushu and the Ryukyu Islands, which are also being primarily divided by their islands (Nakajima & Masuda, 1985; Suzuki & Okano, 2000; Suzuki & Kawai, 2011). The distribution patterns observed by which different species inhabit each island (i.e., tend to be speciated at the island level) could be a general pattern for freshwater crabs.

Freshwater species that adapted to mountain streams, to which freshwater crabs are adapted, have been reported to be more likely to exhibit genetic regionality (Tominaga et al., 2013, 2016; Takenaka & Tojo, 2019; Takenaka et al., 2019; Tojo et al., 2021). Similarly, freshwater crabs in other regions (e.g., Taiwan, China, Africa) have been reported to be genetically differentiated on a regional basis (Daniels et al., 2006; Shih et al., 2006). However, as a result of our genetic analysis of *G. dehaani*, no genetic regionality was detected within each island (however, in the large islands such as Honshu, Shikoku and Kyushu, several intra-islandic genetic lineages were detected). Since adult *G. dehaani* has some capability for terrestrial movement, it was considered possible that gene flow could occur over a wide area within larger scale islands.

Regarding the inconsistency in the genetic clade of the Sado Island population, the Sado specimen was positioned in Clade 1 based on genetic analysis of the mtDNA COI region; however, the Sado specimen was positioned in Clade 2 based on genetic analysis of the combined datasets. If this inconsistency in the genetic positioning of the Sado Island population is correct, the Sado Island population may be derived from an introgression between ancestor populations of Clade 1 observed in Honshu and of Clade 2 observed in western Japan and the Oki Islands. However, only a single sample from Sado Island could be analyzed. Regarding the cladogram based on the mtDNA COI region, it was composed of short branches except for haplotype H36 detected from Sado Island, and only H36 was observed to be genetically differentiated from the other haplotypes with long branches. The cladogram based on the combined dataset indicates genetic differentiation between the genotype G14 detected from Sado Island and other genotypes. Therefore, since there is the possibility of an error due to the single sample, it is necessary to further carefully investigate the phylogenetic relationship of the Sado Island population. Clades 6 and 10 also were based on single samples, so further investigation is also needed, but there is no doubt that they are genetically differentiated from the other clades.

### 4.2 Unexpected distribution expansion: dispersal via two different routes

We have shown *G. dehaani* to be genetically differentiated between islands, having migrated northward terrestrially, but one exceptional clade was detected. That is, Clade 4, detected on the Izu Peninsula, the Miura Peninsula, and the Boso Peninsula. These specimens were all closely related to the clade detected in Kyushu, Shikoku and the Northern Ryukyu Islands (Clades 3, 5–7). As for the results of the ABC analysis, Clade 4 was evaluated to be monophyletic with Clades 5 and 6, and these clades were also evaluated as monophyletic with Clade 3. This means that the Clade 4 shows a disjunct distribution that is closely related to the clades that are widely geographically separated across Honshu and Shikoku. Also, the Izu, Miura and Boso Peninsulas are connected by the Honshu landmass; however, Clade 4 was only detected on these three peninsula areas (Clade 4 was not detected in the inland areas of Honshu), and as such, these populations are isolated from each other. From this distribution pattern, it is considered that an ancestor lineage of Clade 4 migrated from Kyushu or the Northern Ryukyu Islands via ocean currents (Fig. 5). However, since *G. dehaani* lives in the upper regions of rivers for its entire life unlike many crustaceans (Nakajima & Masuda, 1985), this result was unexpected. Shokita (1996) and Yea et al. (2007) also reported that *G. dehaani* cannot survive in seawater. Even so, it was unclear whether or not they could survive for a number of days in the ocean. In order to investigate the potential for ocean current dispersion, we experimentally evaluated the seawater tolerance of *G. dehaani*. This result was surprising and contrary to our initial expectations. In our seawater tolerance experiments, *G. dehaani* survived for at least 15 days in seawater environments. In fact, previous study showed that two freshwater crab families (Potamidae and Gecarcinucidae) can survive in sea water (Esser & Cumberlidge, 2011).

**Figure 5.**
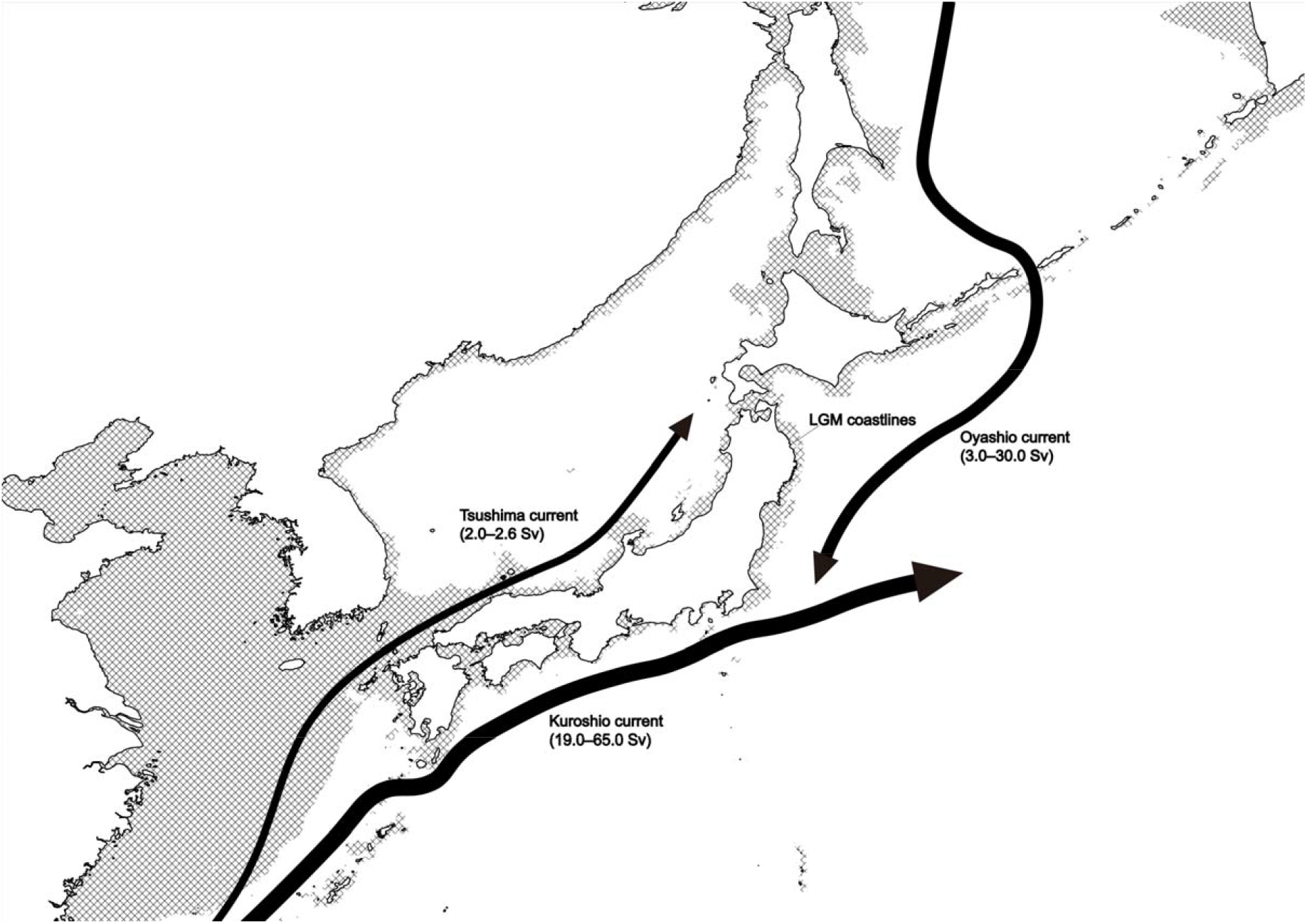
LGM coastlines and major currents around the Japanese Islands. LGM coastlines (Gent et al., 2011), and Sverdrup (Sv: a non-SI metric unit of flow) value of each ocean current are indicated based on previous research (Takikawa, 2005; Andres et al., 2015; Qiu, 2019).

In addition, we determined that *G. dehaani* survived when returned to freshwater after being reared in seawater for 15 days. That is, it was shown that from a physiological and ecological viewpoint that oceanic dispersal of *G. dehaani* is a possibility. Seawater tolerance for as long as 15 days provides the potential for migration via ocean currents that could carry this crab a considerable distance. It was also shown that it could be possible for *G. dehaani* to subsequently survive in freshwater areas after migrating to the Izu Peninsula via ocean currents. Although some cases of seawater crabs migrating via ocean currents have been reported, cases of pure freshwater crabs doing so are extremely rare (Shih et al., 2004; Daniels, S. R. 2011; Jesse et al., 2009; Esser & Cumberlidge, 2011). Considering the geographical genetic structure of *G. dehaani* in this study, the most likely interpretation is that that ocean current dispersion has occurred.

In addition, Clade 2 detected from Suo-Oshima Island, western regions of Honshu and Sado Island was also suggested to migrate via ocean currents. Clade 2 may have migrated via an ocean current on the Sea of Japan side, compared to Clade 4 which would have migrated via Pacific Ocean currents (Fig. 5). There are strong ocean currents (i.e., the “Kuroshio” currents) southwest to northwest on the Pacific Ocean side and the Sea of Japan side, and these conclusions are consistent with these current flow paths (Fig. 5).

In this way, these dispersion events across dispersion barriers for such species by random and/or abnormal events, such as typhoons, hurricanes, floods, changes in the flow routes of ocean currents, and wind direction, are known as “Sweepstakes” events, as suggested by Simpson (1940). For example, the distribution of mammals, amphibians and reptiles that could not cross the sea in Madagascar were explained as “sweepstakes” events (Stankiewicz et al., 2006; Katinas et al., 2013). For *G. dehaani*, almost all clades and species are separated by sea regions, whereby only a few had any potential to be dispersed by ocean currents. Therefore, the existence of parapatric distributions with the clade dispersed on the mainland of Honshu is interesting.

Japanese organisms originating from the southern and western regions migrated to the Japanese Islands via land bridges and expanded their distribution areas across land in a northward direction (Motokawa & Kajihara, 2017; Tojo et al., 2017). Also, it has been reported that species adapted to environments along coasts and organisms that spend a part or all of their lives in the sea rode ocean currents and thereby expanded their distribution areas (Niikura et al., 2015; Abdullah et al., 2017; Ueno et al., 2020; Nakahama et al., 2022). The dispersion routes and methods may be dependent on the environments that they are adapted and/or ecological features. Under such circumstances, it was suggested that *G. dehaani* dispersed via land and ocean toward northern regions via dual routes that differed greatly depending on the system. To date, cases of such dual route species dispersal across both land and ocean have not been reported. We have identified a group that has expanded its distribution area using these dual heterogeneous routes. The evolutionary history of *G. dehaani* estimated in this study is considered to be as follows. The ancestral lineage of *G. dehaani* firstly distributed in a southwestern region and then migrated northward via land bridges. After this expansion of its distribution area within Honshu, the ancestral lineage of Clade 4 dispersed from southern Kyushu and the Northern Ryukyu Islands to the Izu Peninsula, the Miura Peninsula and the Boso Peninsula via ocean currents. As a result, multiple clades of different origins were established within Honshu.

Similarly, within the taxa that are thought to have migrated via the Pacific Ocean, some clades of freshwater shrimps of the *Palaemon* genus distributed in the southern Kyushu region and were also found to have distributed to the Izu Peninsula, the Kii Peninsula and the Muroto Peninsula (Cho et al., 2019). However, some questions remain: “Why was only Clade 4 observed to have migrated via ocean currents?”, and “Why was Clade 4 detected on only three peninsulas?”. What these three peninsulas have in common is that they were once independent islands from Honshu (Shiba, 2016; Takahashi, 2019; Kikuchi, 1997). From a population genetics point of view, it is difficult to change the frequency of the gene pool of an already established population by way of a small number of outside individuals being dispersed to such already established populations. Therefore, in the case of former islands which were isolated from Honshu, freshwater crabs were previously absent, so freshwater crabs successfully dispersed to such islands and could settle into the islands and establish populations. Of course, it cannot be denied that the crabs that make up Clade 1 may have dispersed across the strait to Izu and the other two peninsulas (currently the Izu, Miura and Boso Peninsula), which were once independent islands. However, it would have been difficult to disperse due to the strong ocean current, the “Kuroshio”. In addition, such dispersion from Clade 1 is unlikely from the genetic structures of these peninsula populations. From these viewpoints, the possibility that the crabs in Clade 1 were dispersed after being connected to the land as a peninsula should also rejected. Therefore, it is highly possible that freshwater crabs settled in their optimal niches on as yet unpopulated islands, i.e., the proto Izu Peninsula Island. After that, the population of the Izu Peninsula probably made “stepping stone” migrations, dispersing on to the former islands of the Miura Peninsula and then the Boso Peninsula, which are located further northeast.

### 4.3 Finding several cryptic species

Genetic analyses have often revealed the existence of cryptic species (Pereira-da-Conceicoa et al., 2012; Vuataz et al., 2013; Rutschmann et al., 2014, 2017; Bisconti et al., 2016, 2018; Saito et al., 2018; Yano et al., 2020; Takenaka et al., 2021). In this study, it was revealed that the scale of genetic differentiation observed between the populations of Honshu, Shikoku, and Kyushu, which were morphologically identified as *G. dehaani*, at an interspecific level or greater. In particular, Clades 8 and 9 detected in Shikoku and Kyushu have a larger genetic distance (*p*-distance) than that between other clades. Although the monophyly of Clades 1–7 was highly supported, the monophyly of Clades 1–8 was not so strongly supported. When including Clade 9, Clades 1–9 was not evaluated as being monophyletic. It is necessary to investigate morphological characteristics in more detail, but at least two cryptic species were found on Shikoku and Kyushu in this study.

Besides the Ryukyu Islands, freshwater crabs also inhabit various other islands within the Japanese Islands, and these may contain even more cryptic species. For example, regarding *Geothelphusa sakamotoana* that was treated as one of the outgroups in this study, the H81 haplotype detected on Takara Island of the Tokara Islands is largely genetically differentiated from the haplotypes (H93–99) of *G. sakamotoana* detected on Okinawa-jima and Amami-Oshima Islands, although their phylogenetic relationship is not yet certain. It was also revealed that H93–97 detected in Okinawa Island and H98–99 detected in Amami-Oshima Island are largely genetically differentiated. Also, Clade 6 detected on Tanegashima Island and Clade 10 detected on Kyushu (only from site No.95) were genetically differentiated from the other clades, therefore they come from isolated genetic lineages limited to each geographical area.

Differences in the body color of *G. dehaani* have often been reported in many regions (Chokki, 1976, 1980; Nakajima & Masuda, 1985; Suzuki & Tsuda, 1991), but each of these previous studies focused on a restricted regional scale, so the details of the relationships between body color and phylogeny are unknown. Although a red body color and blue body color specimens of *G. dehaani* were collected within the same stream on Shikoku in this study, no detected genetic differences were found between each body color type (Fig. 1D, E: st. 116). This suggests that there is basically no relationship between body color and genetic differences in *G. dehaani* (H82 and H83: Table 1; Fig. 3). Previously, Aotsuka et al. (1995) reported two body colors (i.e., dark brown and blue) of *G. dehaani* that exhibited some degree of genetic differentiation using isozyme variation analysis. The population analyzed in a previous study (Aotsuka et al., 1995) constituted Clade 1 in the present genetic analyses results. Meanwhile, genetically differentiated crabs consisting of Clade 4 inhabited an area close to this site (Aotsuka et al., 1995), and the body color of the freshwater crabs in Clade 4 was pale bluish (this colors treated to be as “blue” color in previous studies). Almost all samples collected from the Izu Peninsula (Clade 4: Fig. 1F) and the Northern Ryukyu Islands (Clades 3 and 6) had blue bodies. Therefore, the contradiction between the previous study and our results may be explained by the possibility of the existence of hybrids in Clades 1 and 4. Aotsuka et al. (1995) conducted isozyme analysis, and found the existence of more polymorphisms than the genetic loci in our analyses, so it is possible that the results of the previous study may reflect more recent events. Therefore, it is highly possible that Clade 4 made a secondary contact, then hybridized with Clade 1 after migrating to the Izu Peninsula via ocean currents. To establish if this is the case, it is necessary to analyze more sensitive molecular markers (SNPs, SSR, etc.). It is not known whether the blue body color occurred following migration via ocean currents or as a result of the “founder effect”. It will be an interesting to investigate any correlations between body color differences and phylogenetic relationships.

In conclusion, it is known that the species distributed in the Ryukyu Islands are primarily differentiated by straits, and it was also found that species or genetic clades distributed in Honshu, Shikoku, and Kyushu are also fundamentally differentiated by straits. Therefore, it can be generally said that freshwater crab dispersal and/or connectivity between populations is restricted by straits and that speciation occurs between isolated populations.

However, strong evidence for dispersion via ocean currents was detected (i.e., a “sweepstake”), and it was also determined that *G. dehaani* could survive in seawater. At first glance, the results were contradictory, but since *G. dehaani* inhabits the upper reaches of river systems, it is thought that dispersion events via ocean currents are rare. That is, ocean current-based dispersal is unlikely to occur in habitats of large river systems. However, in small-scale river systems such as on small-scale islands and peninsulas, it is often the case that upstream-inhabiting freshwater crabs are swept away to sea areas during floods. As a result, it is thought that ocean current-based dispersal has occurred to such islands and/or peninsula habitats. In addition, even if freshwater crabs succeed in dispersal via ocean currents, it is unlikely that they can successfully colonize habitats if other lineage(s) are already distributed there, so dispersion events via ocean currents are considered to be an extremely rare phenomenon. In fact, Clade 4 detected on the Izu Peninsula, the Miura Peninsula and the Boso Peninsula was found to have become significantly genetically differentiated from the other clades.

The highlights of this study were the discovery of several cryptic species/lineages or undescribed species, and the completely different heterogeneous dual dispersal pathways detected within a single species; i.e., both land and ocean routes. This is the first observation of such a unique mode of expansion of a species’ distribution area, providing new knowledge and insight regarding the development of biodiversity in Japanese organisms.

## Acknowledgements

We express our thanks to the Office of Numazu River and National Highway (Chubu Regional Development Bureau, Ministry of Land Infrastructure and Transport) and Dr. Sekiné, K. (Risho University), Dr. Kunishima T. (Wakayama Prefectural Museum of Natural History), Mr. Takata K. (Kyoto University), Dr. Toyoda K. (Niigata University), the members of Tojo Laboratory (Shinshu University); Dr. Suzuki T., Dr. Komaki S., Dr. Tamura Mr., Kogawara H., Dr. Okamoto S., Mr. Ueki G., Ms. Koike K., Mr. Fujimori T., Mr. Ozaki T., Ms. Momose K., Mr. Inoue K., Mr. Tomizawa R., Mr. Yoshida M., Mr. Yoshida T. This study was supported by grants from the Pro-Natura Foundation Japan (2016-2017, KT) and the River Environment Fund (25-1215-016, 2017-5211-025) of River and Watershed Environment Management, by Muroto UNESCO Global Geopark Research Grant 2018, 2020 (M.T.), by Oki Islands UNESCO Global Geopark Research Grant 2019 (K.Y.), by Kuroshio Biological Research Foundation Research Grant 2019 (K.Y.), and by a research grant from the Institute of Mountain Science, Shinshu University (K.T.).

## Biosketches

**Masaki Takenaka** is interested in the biogeography of inland water organisms in Asia, with special emphasis on mayflies. He is an Associate Professor at the Department of Biology, Faculty of Science, Shinshu University.

**Koki Yano** received a Ph.D. from Shinshu University for his study on the phylogeny and evolutional ecology of aquatic insects. He is now working as part of a Research Fellowship for Young Scientists of JSPS at the National Institute for Basic Biology. His main interest is the biogeography of cosmopolitan freshwater organisms.

**Koji Tojo** is a Professor in the Department of Biology, Faculty of Science, Shinshu University. He is interested in the biogeography of freshwater animals on a worldwide scale. Members of his laboratory are engaged in research on biogeography and the evolutionary ecology of freshwater animals.

## Author contributions

M.T., K.Y. and K.T. designed, managed the study, and performed sample collection; M.T. and K.Y. mainly performed laboratory work and phylogenetic analyses; M.T., K.Y. and K.T. wrote and reviewed the manuscript.

---

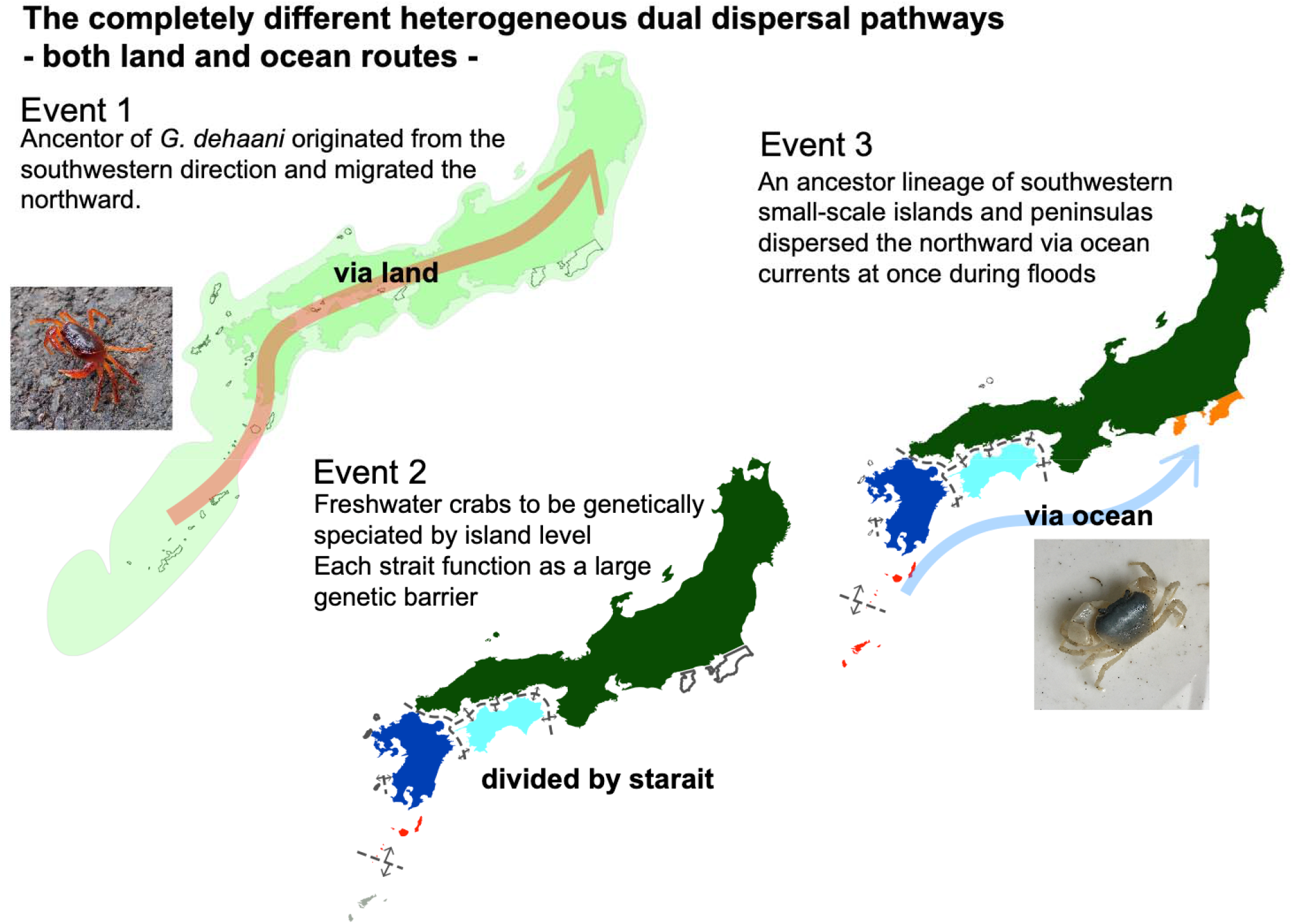

## References

Abdullah, M. F., Cheng, J. H., Chen, T. I., & Imai, H. (2017). Development of compound polymorphic microsatellite markers for the pronghorn spiny lobster *Panulirus penicillatus* and comparison of microsatellite data with those of a previous mitochondrial DNA study performed in the northwestern Pacific. Biogeography, 19, 61–68.

Andres, M., Jan S., Sanford T. B., Mensah V., Centurioni L. R., & Book, J. W. (2015). Mean structure and variability of the Kuroshio from northeastern Taiwan to southwestern Japan. Oceanography, 28(4), 84–95.

Aotsuka, T., Suzuki, T., Moriya, T., & Inaba, A. (1995). Genetic differentiation in Japanese freshwater crab, *Geothelphusa dehaani* (White): isozyme variation among natural populations in Kanagawa Prefecture and Tokyo. Zoological Science, 12(4), 427–434.

Avise, J. C. (2000). Phylogeography: the history and formation of species. Massachusetts, Harvard University press,

Banks, S. C., Ling, S. D., Johnson, C. R., Piggott, M. P., Williamson, J. E., & Beheregaray, L. B. (2010). Genetic structure of a recent climate change-driven range extension. Molecular Ecology, 19(10), 2011–2024.

Bisconti, R., Canestrelli, D., Tenchini, R., Belfiore, C., Buffagni, A., & Nascetti, G. (2016). Cryptic diversity and multiple origins of the widespread mayfly species group *Baetis rhodani* (Ephemeroptera: Baetidae) on northwestern Mediterranean islands. Ecology and Evolution, 6(21), 7901–7910.

Bisconti, R., Tenchini, R., Belfiore, C., Nascetti, G., & Canestrelli, D. (2018). Fast and accurate identification of cryptic and sympatric mayfly species of the *Baetis rhodani* group. BMC Research Notes, 11(1), 1–6.

Chokki, H. (1976). Preliminary report of the colouration of freshwater crab, *Geothelphusa dehaani* (White), with special reference to its distribution. Researches on Crustacea (Former issue of Crustacean Research), 7, 1–13.

Chokki, H. (1980). Notes on the colouration of freshwater crab, *Geothelphusa dehaani* (White), in northern districts of Japan. Researches on Crustacea, 10, 57–60.

Chow, S., Yanagimoto, T., Konishi, K., Ichikawa, T., Komatsu, N., Maruyama, T., Ikeda, M., Nohara, K., Oonuki, T., & Imai, T. (2019). Genetic diversity in type B of Palaemon paucidens. Aquatic Animals AA2019-11.

Cook, B. D., Pringle, C. M., & Hughes, J. M. (2008). Phylogeography of an island endemic, the Puerto Rican freshwater crab (Epilobocera sinuatifrons). Journal of Heredity, 99(2), 157–164.

Copilaş-Ciocianu, D., & Petrusek, A. (2017). Phylogeography of a freshwater crustacean species complex reflects a long-gone archipelago. Journal of Biogeography, 44(2), 421–432.

Cornuet, J. M., Pudlo, P., Veyssier, J., Dehne-Garcia, A., Gautier, M., Leblois, R., Marin, J. M., & Estoup, A. (2014). DIYABC v2. 0: a software to make approximate Bayesian computation inferences about population history using single nucleotide polymorphism, DNA sequence and microsatellite data. Bioinformatics, 30(8), 1187–1189.

Daniels, S. R., Stewart, B. A., Gouws, G., Cunningham, M., & Matthee, C. A. (2002). Phylogenetic relationships of the southern African freshwater crab fauna (Decapoda: Potamonautidae: *Potamonautes)* derived from multiple data sets reveal biogeographic patterning. Molecular Phylogenetics and Evolution, 25(3), 511–523.

Daniels, S. R., Gouws, G., Stewart, B. A., & Coke, M. (2003). Molecular and morphometric data demonstrate the presence of cryptic lineages among freshwater crabs (Decapoda: Potamonautidae: *Potamonautes)* from the Drakensberg Mountains, South Africa. Biological Journal of the Linnean Society, 78(1), 129–147.

Daniels, S. R., Gouws, G., & Crandall, K. A. (2006). Phylogeographic patterning in a freshwater crab species (Decapoda: Potamonautidae: *Potamonautes)* reveals the signature of historical climatic oscillations. Journal of Biogeography, 33(9), 1538–1549.

Daniels, S. R. (2011). Reconstructing the colonisation and diversification history of the endemic freshwater crab *(Seychellum alluaudi)* in the granitic and volcanic Seychelles Archipelago. Molecular Phylogenetics and Evolution, 61(2), 534–542.

Daniels, S. R., Phiri, E. E., Klaus, S., Albrecht, C., & Cumberlidge, N. (2015). Multilocus phylogeny of the Afrotropical freshwater crab fauna reveals historical drainage connectivity and transoceanic dispersal since the Eocene. Systematic Biology, 64(4), 549–567.

Esser, L. J., & Cumberlidge, N. (2011). Evidence that salt water may not be a barrier to the dispersal of Asian freshwater crabs. The Raffles Bulletin of Zoology, 59(2), 259–268.

Felsenstein, J. (1981). Evolutionary trees from DNA sequences: a maximum likelihood approach. Journal of molecular evolution, 17(6), 368–376.

Furuya Y, & Yamaoka J. (2017) The mystery of Freshwater crab “blue”. Kochi, Minami-no-kaze.

García-Verdugo, C., Mairal, M., Monroy, P., Sajeva, M., & Caujapé-Castells, J. (2017). The loss of dispersal on islands hypothesis revisited: Implementing phylogeography to investigate evolution of dispersal traits in *Periploca* (Apocynaceae). Journal of Biogeography, 44(11), 2595–2606.

Gent, P. R., Danabasoglu, G., Donner, L. J., Holland, M. M., Hunke, E. C., Jayne, S. R., Lawrence, D. M., Neale, R. B., Rasch, P. J., Vertenstein, M., Worley, P. H., Yang, Z. L., & Zhang, M. (2011). The community climate system model version 4. Journal of Climate, 24(19), 4973–4991.

Huelsenbeck, J. P., & Ronquist, F. (2001). MRBAYES: Bayesian inference of phylogenetic trees. Bioinformatics, 17(8), 754–755.

Ikeda, M., Suzuki, T., & Fujio, Y. (1998). Genetic differentiation among populations of Japanese freshwater crab, *Geotheiphusa dehaani* (White), with Reference to the Body Color Variation. Benthos Research, 53(1), 47–52.

Jesse, R., Pfenninger, M., Fratini, S., Scalici, M., Streit, B., & Schubart, C. D. (2009). Disjunct distribution of the Mediterranean freshwater crab *Potamon fluviatile—*natural expansion or human introduction?. Biological Invasions, 11(10), 2209–2221.

Katinas, L., Crisci, J. V., Hoch, P., Tellería, M. C., & Apodaca, M. J. (2013). Trans-oceanic dispersal and evolution of early composites (Asteraceae). Perspectives in Plant Ecology, Evolution and Systematics, 15(5), 269–280.

Kikuchi, T. (1997). From proto-Tokyo Bay to Tokyo Bay: The role of structural movements in the transitional period (special feature “Tokyo Bay”). Chigakukyoiku and Kagaku-undo, 28, 1–10.

Kobayashi, S. (2000). Distribution pattern and ecology of brachyuran crabs in the riverine environment their significance in the ecosystem and present condition. Ecology and Civil Engineering, 3(1), 113–130.

Koizumi, I., Usio, N., Kawai, T., Azuma, N., & Masuda, R. (2012). Loss of genetic diversity means loss of geological information: the endangered Japanese crayfish exhibits remarkable historical footprints. PLOS ONE, 7(3), e33986.

McConaugha, J. R. (1992). Decapod larvae: dispersal, mortality, and ecology. A working hypothesis. American Zoologist, 32(3), 512–523.

Miyake S. (1983). Japanese crustacean decapods and stomatopods in color v2. Tokyo, Hoikusha.

Motokawa, M., & Kajihara, H. (2017). Species diversity of animals in Japan. Tokyo, Springer Japan.

Nakahama, N., Okano, R., Nishimoto, Y., Matsuo, A., Ito, N., & Suyama, Y. (2022). Possible dispersal of the coastal and subterranean carabid beetle *Thalassoduvalius masidai* (Coleoptera) by ocean currents. Biological Journal of the Linnean Society, 135(2), 265–276.

Nakajima, K., & Masuda, T. (1985). Identification of local populations of freshwater crab *Geothelphusa dehaani* (White). Bulletin of the Japanese Society of Scientific Fisheries, 51(2), 175–181.

Niikura, M., Honda, M., & Yahata, K. (2015). Phylogeography of semiterrestrial isopod, *Tylos granuliferus*, on East Asian coasts. Zoological Science, 32(1), 105–113.

Okano, T., Suzuki, H., Hiwatashi, Y., Nagoshi, F., & Miura, T. (2000). Genetic divergence among local populations of the Japanese freshwater crab *Geothelphusa dehaani* (Decapoda, Brachyura, Potamidae) from southern Kyushu, Japan. Journal of Crustacean Biology, 20(4), 759–768.

Orsini, L., Vanoverbeke, J., Swillen, I., Mergeay, J., & De Meester, L. (2013). Drivers of population genetic differentiation in the wild: isolation by dispersal limitation, isolation by adaptation and isolation by colonization. Molecular Ecology, 22(24), 5983–5999.

Page, T. J., Cook, B. D., von Rintelen, T., von Rintelen, K., & Hughes, J. M. (2008). Evolutionary relationships of atyid shrimps imply both ancient Caribbean radiations and common marine dispersals. Journal of the North American Benthological Society, 27(1), 68–83.

Pearse, A. S. (1927). The migration of animals from the ocean into freshwater and land habitats. The American Naturalist, 61(676), 466–476.

Pereira-da-Conceicoa, L. L., Price, B. W., Barber-James, H. M., Barker, N. P., De Moor, F. C., & Villet, M. H. (2012). Cryptic variation in an ecological indicator organism: mitochondrial and nuclear DNA sequence data confirm distinct lineages of *Baetis harrisoni* Barnard (Ephemeroptera: Baetidae) in southern Africa. BMC Evolutionary Biology, 12(1), 1–14.

Promnun, P., Tandavanitj, N., Kongrit, C., Kongsatree, K., Kongpraphan, P., Dongkumfu, W., Kumsuan, D., & Khudamrongsawat, J. (2021). Phylogeography and ecological niche modeling reveal evolutionary history of *Leiolepis ocellata* (Squamata, Leiolepidae). Ecology and Evolution, 11(5), 2221–2233.

Qiu, B. (2019). Kuroshio and Oyashio currents. In J. H. Steele, S. A. Thorpe, & K. K. Turekian (Eds.), Encyclopedia of ocean sciences. (pp. 1413–1425). New York, Academic Press.

Rambaut, A. (2009). FigTree, Version 1.3.1. Available at: http://tree.bio.ed.ac.uk/sofiware/figtree/.

Rambaut, A., & Drummond, A. J. (2007). Tracer, Version 1.5. MCMC Trace Analysis Tool. Available at: http://tree.bio.ed.ac.uk/software/tracer/.

Rocha, L. A., Rocha, C. R., Robertson, D. R., & Bowen, B. W. (2008). Comparative phylogeography of Atlantic reef fishes indicates both origin and accumulation of diversity in the Caribbean. BMC Evolutionary Biology, 8(1), 1–16.

Rutschmann, S., Gattolliat, J. L., Hughes, S. J., Báez, M., Sartori, M., & Monaghan, M. T. (2014). Evolution and island endemism of morphologically cryptic *Baetis* and *Cloeon* species (Ephemeroptera, Baetidae) on the Canary Islands and Madeira. Freshwater Biology, 59(12), 2516–2527.

Rutschmann, S., Detering, H., Simon, S., Funk, D. H., Gattolliat, J. L., Hughes, S. J., Raposeiro, P. M., DeSalle, R., Sartori, M., & Monaghan, M. T. (2017). Colonization and diversification of aquatic insects on three Macaronesian archipelagos using 59 nuclear loci derived from a draft genome. Molecular Phylogenetics and Evolution, 107, 27–38.

Saito, R., Kato, S., Kuranishi, R. B., Nozaki, T., Fujino, T., & Tojo, K. (2018). Phylogeographic analyses of the *Stenopsyche* caddisflies (Trichoptera: Stenopsychidae) of the Asian Region. Freshwater Science, 37(3), 562–572.

Schwarz, G. (1978). Estimating the dimension of a model. The Annals of Statistics, 6(2) 461–464.

Shiba, M. (2016) Did Izu Peninsula come from south? Journal of Fossil Research, 49, 35–43.

Shih, H. T., & Ng, P. K. (2011). Diversity and biogeography of freshwater crabs (Crustacea: Brachyura: Potamidae, Gecarcinucidae) from east Asia. Systematics and Biodiversity, 9(1), 1–16.

Shih, H. T., Ng, P. K., & Chang, H. W. (2004). Systematics of the genus *Geothelphusa* (Crustacea, Decapoda, Brachyura, Potamidae) from southern Taiwan: a molecular appraisal. Zoological Studies 43(3), 561–570.

Shih, H. T., Hung, H. C., Schubart, C. D., Chen, C. A., & Chang, H. W. (2006). Intraspecific genetic diversity of the endemic freshwater crab *Candidiopotamon rathbunae* (Decapoda, Brachyura, Potamidae) reflects five million years of the geological history of Taiwan. Journal of Biogeography, 33(6), 980–989.

Shih, H. T., Ng, P. K., Schubart, C. D., & Chang, H. W. (2007). Phylogeny and phylogeography of the genus *Geothelphusa* (Crustacea: Decapoda, Brachyura, Potamidae) in southwestern Taiwan based on two mitochondrial genes. Zoological Science, 24(1), 57–66.

Shih, H. T., Zhou, X. M., Chen, G. X., Chien, I. C., & Ng, P. K. (2011). Recent vicariant and dispersal events affecting the phylogeny and biogeography of East Asian freshwater crab genus *Nanhaipotamon* (Decapoda: Potamidae). Molecular Phylogenetics and Evolution, 58(3), 427–438.

Shokita, S. (1996). The origin of land-locked freshwater shrimps and potamoids from the Ryukyu Islands, southern Japan. Journal of Geography (Chigaku Zasshi), 105(3), 343–353.

Shokita, S., Naruse, T., & Fujii, H. (2002). *Geothelphusa miyakoensis*, new species of freshwater crab (Crustacea: Decapoda: Brachyura: Potamidae) from Miyako Island, Southern Ryukyus, Japan. Raffles Bulletin of Zoology, 50(2), 443–448.

Sota, T., & Vogler, A. P. (2001). Incongruence of mitochondrial and nuclear gene trees in the carabid beetles *Ohomopterus*. Systematic Biology, 50(1), 39–59.

Stamatakis, A. (2014). RAxML version 8: a tool for phylogenetic analysis and post-analysis of large phylogenies. Bioinformatics, 30(9), 1312–1313.

Stankiewicz, J., Thiart, C., Masters, J. C., & De Wit, M. J. (2006). Did lemurs have sweepstake tickets? An exploration of Simpson’s model for the colonization of Madagascar by mammals. Journal of Biogeography, 33(2), 221–235.

Su, Z. H., Ohama, T., Okada, T. S., Nakamura, K., Ishikawa, R., & Osawa, S. (1996). Geography-linked phylogeny of the *Damaster* ground beetles inferred from mitochondrial ND5 gene sequences. Journal of Molecular Evolution, 42(2), 130–134.

Su, Z. H., Tominaga, O., Okamoto, M., & Osawa, S. (1998). Origin and diversification of hindwingless *Damaster* ground beetles within the Japanese islands as deduced from mitochondrial ND5 gene sequences (Coleoptera, Carabidae). Molecular Biology and Evolution, 15(8), 1026–1039.

Suzuki, T. (1992). Coloration and distribution of the Japanese freshwater crab, *Geothelphusa dehaani* (White), in the Hanamizu River on Tanzawa Mountains, Kanagawa Prefecture. Natural History Report of Kanagawa, 13, 55–64.

Suzuki, H. (1994). A new freshwater crab of the genus *Geothelphusa* (Crustacea: Decapoda: Brachyura: Potamidae) from Kagoshima Prefecture, southern Kyushu, Japan. Proceedings of the Biological Society of Washington, 107, 318–324.

Suzuki, H., & Kawai, T. (2011). Two new freshwater crabs of the genus *Geothelphusa* Stimpson, 1858 (Crustacea: Decapoda: Brachyura: Potamidae) from islands of southern Kyushu, Japan. Crustacean Research, 40, 21–31.

Suzuki, H., & Naruse, T., (2012). Freshwater Decapoda Crustaceans in Japan. In T. Kawai & K. Nakata (Eds.), Shrimps, Crabs and Crawfish—Conservation and Biology of Freshwater Crustaceans—. (pp. 39–73). Tokyo, Seibutsu-kenkyusha.

Suzuki, H., & Okano, T. (2000). A new freshwater crab of the genus *Geothelphusa* Stimpson, 1858 (Crustacea: Decapoda: Brachyura: Potamidae) from Yakushima Island, southern Kyushu, Japan. Proceedings of the Biological Society of Washington, 113(1), 30–38.

Suzuki, H., & Tsuda, E. (1991). Study on the color variation and distribution of a freshwater crab, *Geothelphusa dehaani* (White), in Kagoshima Prefecture. Benthos Research, 1991(41), 37–46.

Takahashi, M. (2019) Geology of the Kanto district as recorded of back-arc spreading of the Japan Sea. Journal of Fossil Research, 52, 1–10.

Takenaka, M., & Tojo, K. (2019). Ancient origin of a dipteromimid mayfly family endemic to the Japanese Islands and its genetic differentiation across tectonic faults. Biological Journal of the Linnean Society, 126(3), 555–573.

Takenaka, M., Tokiwa, T., & Tojo, K. (2019). Concordance between molecular biogeography of *Dipteromimus tipuliformis* and geological history in the local fine scale (Ephemeroptera, Dipteromimidae). Molecular Phylogenetics and Evolution, 139, 106547.

Takenaka, M., Yano, K., Suzuki, T., & Tojo, K. (2021). Development of novel PCR primer sets for DNA metabarcoding of aquatic insects, and the discovery of some cryptic species. bioRxiv.

Takikawa, T. (2005). The Tsushima Warm Current trough Tsushima Straits estimated from ferryboat ADCP data. Journal of Physical Oceanography, 35(6), 1154–1168.

Tanabe, A. S. (2007). Kakusan: a computer program to automate the selection of a nucleotide substitution model and the configuration of a mixed model on multilocus data. Molecular Ecology Notes, 7(6), 962–964.

Tanabe, A. S. (2008). MrBayes5D. Software distributed by the author at http://www.fifthdimension.jp/

Tojo, K., Sekiné, K., Takenaka, M., Isaka, Y., Komaki, S., Suzuki, T., & Schoville, S. D. (2017). Species diversity of insects in Japan: their origins and diversification processes. Entomological Science, 20(1), 357–381.

Tojo, K., Miyairi, K., Kato, Y., Sakano, A., & Suzuki, T. (2021). A description of the second species of the genus *Bleptus* Eaton, 1885 (Ephemeroptera: Heptageniidae) from Japan, and phylogenetic relationships of two *Bleptus* mayflies inferred from mitochondrial and nuclear gene sequences. Zootaxa, 4974(2), 333360.

Tominaga, A., Matsui, M., Yoshikawa, N., Nishikawa, K., Hayashi, T., Misawa, Y., Tanabe, S., & Ota, H. (2013). Phylogeny and historical demography of *Cynops pyrrhogaster* (Amphibia: Urodela): taxonomic relationships and distributional changes associated with climatic oscillations. Molecular Phylogenetics and Evolution, 66(3), 654–667.

Tominaga, K., Nakajima, J., & Watanabe, K. (2016). Cryptic divergence and phylogeography of the pike gudgeon *Pseudogobio esocinus* (Teleostei: Cyprinidae): a comprehensive case of freshwater phylogeography in Japan. Ichthyological Research, 63(1), 79–93.

Toyota, K., & Seki, S. (2019). Picture book of Japanese freshwater/brackish water shrimp and crab. Tokyo, Midorishobo.

Ueno, H., Kitagawa, K., & Matsubayashi, K. W. (2020). Unexpectedly long survivorship on seawater of multiple coastal beetles indicates the possibility of “floating dispersal” for transoceanic migration. Entomological Science, 23(3), 294–296.

Van der Stocken, T., Wee, A. K., De Ryck, D. J., Vanschoenwinkel, B., Friess, D. A., Dahdouh-Guebas, F., Simard, M., Koedam, N., & Webb, E. L. (2019). A general framework for propagule dispersal in mangroves. Biological Reviews, 94(4), 1547–1575.

Vuataz, L., Sartori, M., Gattolliat, J. L., & Monaghan, M. T. (2013). Endemism and diversification in freshwater insects of Madagascar revealed by coalescent and phylogenetic analysis of museum and field collections. Molecular Phylogenetics and Evolution, 66(3), 979–91.

Waters, J. M., Emerson, B. C., Arribas, P., & McCulloch, G. A. (2020). Dispersal reduction: causes, genomic mechanisms, and evolutionary consequences. Trends in Ecology & Evolution, 35(6), 512–522.

Wowor, D., Muthu, V., Meier, R., Balke, M., Cai, Y., & Ng, P. K. (2009). Evolution of life history traits in Asian freshwater prawns of the genus *Macrobrachium* (Crustacea: Decapoda: Palaemonidae) based on multilocus molecular phylogenetic analysis. Molecular Phylogenetics and Evolution, 52(2), 340–350.

Yano, K., Takenaka, M., & Tojo, K. (2019). Genealogical position of Japanese populations of the globally distributed mayfly *Cloeon dipterum* and related species (Ephemeroptera, Baetidae): A molecular phylogeographic analysis. Zoological Science, 36(6), 479–489.

Yano, K., Takenaka, M., Mitamura, T., & Tojo, K. (2020). Identifying a “pseudogene” for the mitochondrial DNA COI region of the corixid aquatic insect, *Hesperocorixa distanti* (Heteroptera, Corixidae). Limnology, 21(3), 319–325.

Yeo, D. C. J., Ng, P. K. L., Cumberlidge, N., Magalhães, C., Daniel, S. R., & Campos, M. R. (2008). Global diversity of crabs (Crustacea: Decapoda: Brachyura) in freshwater. Hydrobiologia, 595(1), 275.

York, K. L., Blacket, M. J., & Appleton, B. R. (2008). The Bassian Isthmus and the major ocean currents of southeast Australia influence the phylogeography and population structure of a southern Australian intertidal barnacle *Catomerus polymerus* (Darwin). Molecular Ecology, 17(8), 1948–1961.

Yorisue, T., Iguchi, A., Yasuda, N., Yoshioka, Y., Sato, T., & Fujita, Y. (2020). Evaluating the effect of overharvesting on genetic diversity and genetic population structure of the coconut crab. Scientific Reports, 10(1), 1–9.

